# PKA-Regulated Carbohydrate Production Protects Cells by Altering Cytoplasmic Biophysical Properties

**DOI:** 10.64898/2026.08.29.747980

**Authors:** Marina Kunzi, Lorena Kronig, Martina Bonassera, Pablo A. Gómez-García, Matthias Peter, Karsten Weis, Gabriel E. Neurohr

## Abstract

Proliferating cells maintain their cytoplasmic density within a narrow range but deviate when entering quiescence or experiencing stress, suggesting active regulation. The mechanisms driving these density adjustments and their impact on cellular function remain unclear. Here, we demonstrate that the conserved cAMP-activated protein kinase A (PKA) is a key regulator of cytoplasmic properties. Inactivation of PKA leads to a drastic increase in cytoplasmic dry mass density and reduced diffusion that depends on the environmental stress response (ESR) transcription factors Msn2/4. This change is mediated by the accumulation of glycogen and trehalose, which have opposing effects on intracellular diffusion. Importantly, the accumulation of these carbohydrates confers stress resistance in distinct ways and independently of their roles as energy sources. Our findings highlight the importance of the biophysical properties of the cytoplasm in stress resistance and the role of glycogen and trehalose in regulating these properties.

## Introduction

Cells must withstand different stresses as environmental conditions fluctuate. Signalling pathways that respond to such stresses and induce appropriate adaptations are critical for the survival of all organisms. Different types of stress often converge on common processes, like the conserved eukaryotic integrated stress response, which downregulates translation initiation in response to starvation, oxidative stress or viral infection ^1^. In budding yeast, the environmental stress response (ESR) is activated by many environmental perturbations, including nutrient starvation, heat, osmotic shock, or oxidative stress and confers increased tolerance to various forms of stress ^2,3^. Activation of the ESR leads to a massive transcriptional rewiring that includes downregulation of ribosome biogenesis, changes in metabolism and induction of chaperones and de-toxifying enzymes ^4,5^.

More recently, it has been recognized that apart from broad remodeling of the transcriptome and proteome, cells also alter the biophysical properties of the cytoplasm in response to stress: cytoplasmic crowding increases in response to glucose starvation and heat stress, while it decreases upon amino acid starvation and cell cycle arrest ^6–11^. Importantly, such changes in crowding coincide with altered cell growth and cell function ^12–15^. How cells regulate cytoplasmic density and crowding in response to stress is poorly understood and it is therefore also unclear whether and how cytoplasmic properties contribute to stress survival.

Recent work points toward classical growth regulatory pathways being involved in altering cytoplasmic properties and survival during stress. TOR complex 1 (TORC1) has been identified as a regulator of cytoplasmic properties through its role in regulating ribosome biogenesis and translation ^8,16,17^. Similarly, components of the Ras-cAMP-PKA signalling cascade have been shown to modulate diffusion and promote viability in energy-depleted cells ^18^. The highly conserved PKA signalling pathway regulates growth, metabolism and stress resistance by sensing both intra- and extracellular glucose and responding to heat shock and oxidative stress ^19–21^. PKA signalling is inactivated by these stresses to halt biomass production and proliferation. Inactive PKA also leads to dephosphorylation and nuclear localization of the ESR transcription factors Msn2/4, where they activate transcription of stress-responsive genes ^4,5^. Among the upregulated transcripts are many genes involved in energy production, proteostasis and carbon metabolism.

A major metabolic adaptation downstream of PKA inactivation is the accumulation of the storage carbohydrates glycogen and trehalose ^22,23^. The expression and activity of the corresponding synthesis and degradation enzymes are regulated by PKA and Msn2/4 ^4,5,24,25^. While trehalose and glycogen are classically viewed as long-term metabolic energy stores mobilized during starvation ^26,27^, they have also been implicated in affecting the biophysical properties of the cytoplasm. Trehalose and glycogen have been reported to increase cytoplasmic viscosity upon heat stress and glucose deprivation to ensure invariant reaction rates ^11^. In support of a biophysical role, trehalose is well-known to confer increased resistance to heat shock and to stabilize proteins under denaturing conditions ^28–32^. However, how glycogen and trehalose modulate intracellular organization and cell survival remains poorly understood.

Here, we combine quantitative measurements of density and diffusion to investigate how PKA and the ESR modulate the biophysical properties of the cytoplasm through production of glycogen and trehalose and how these carbohydrates impact viability during stress. Our results show that glycogen and trehalose confer resistance to heat and nutrient stress, but not by serving as energy reserves; rather, they regulate the biophysical state of the cytoplasm in distinct and opposing ways.

## Results

### PKA regulates cytoplasmic density and diffusion via the environmental stress response

To determine how growth and stress signalling affect the biophysical properties of budding yeast cells, we genetically inactivated PKA and measured how this influences cytoplasmic density. Since PKA is essential ^22^, we used a conditional PKA inactivation (PKAi) strain where all three catalytic isoforms, encoded by *TPK1, TPK2* and *TPK3*, were mutated to be sensitive to the ATP analogue 1NM-PP1 (TPKas, ^33^). These mutations allow for temporally controlled and reversible inhibition of PKA through addition of the ATP analogue 1NM-PP1 ^34^. To measure cell density, we employed two independent methods. Firstly, refractive index holotomography allowed us to quantitatively measure the cytoplasmic dry mass density excluding the vacuole ^35^. PKA inhibition led to a drastic increase in cytoplasmic dry mass density of roughly 50% (**Figures 1A and 1B**). Secondly, we used density-gradient centrifugation of the cells in microcapillaries to measure whole-cell buoyant density (**Figure S1A, S1B, S1C**). The buoyant density for asynchronous cultures of WT cells obtained through this method was 1.083 ± 0.002 g/mL and increased to 1.132 ± 0.002 g/mL upon PKA inhibition (**Figure 1C**). Together, these results identify PKA as a key regulator of cell density.

**Figure 1.**
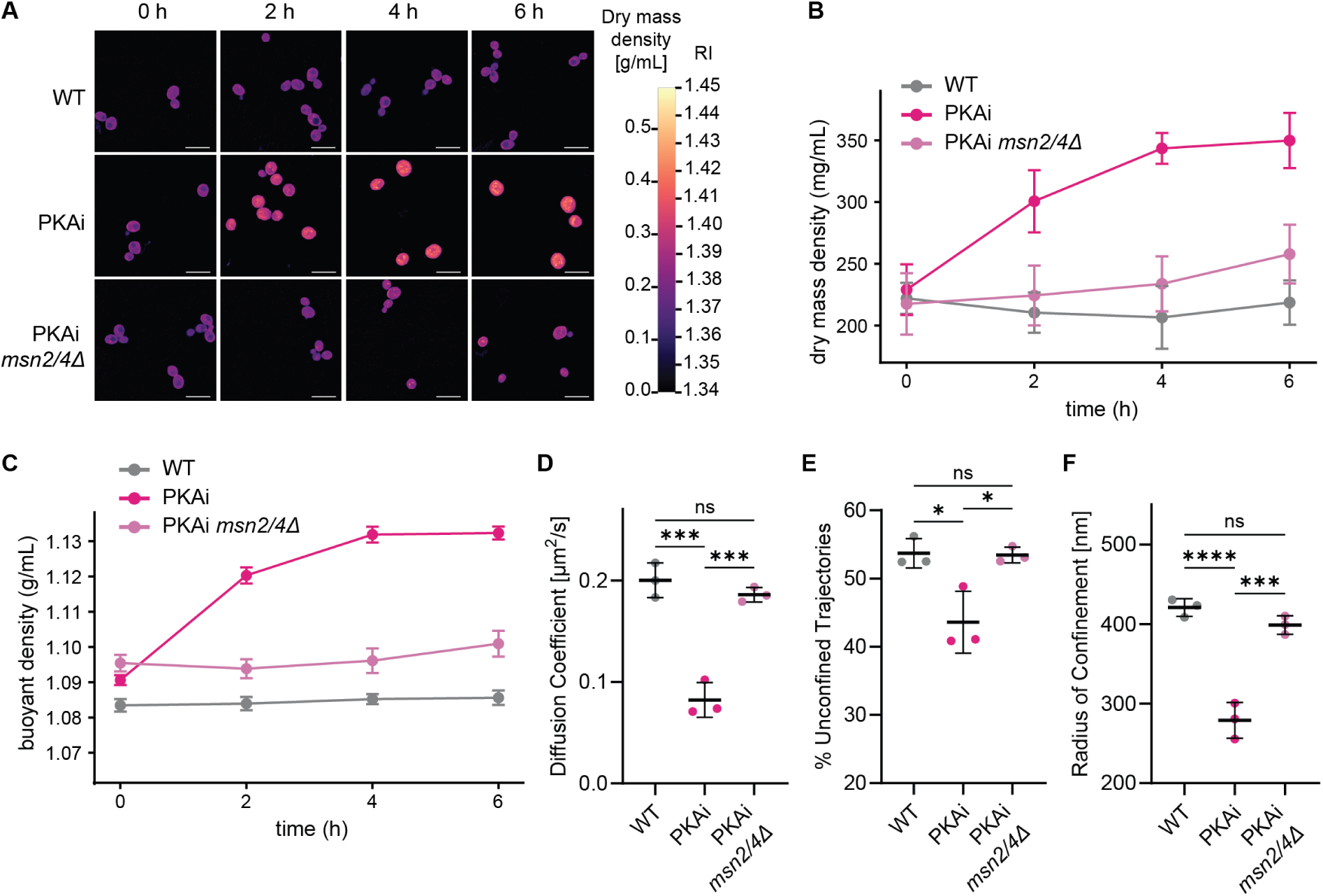
PKA inhibition increases cell density and reduces particle diffusion in an Msn2/4-dependent manner. **(A)** Refractive index holotomography and conversion of refractive index (RI) into dry mass density of wild-type (WT) cells or upon acute PKA-inhibition (PKAi) in otherwise wild-type or *msn2/4Δ* mutant cells. ATP analogue 1NM-PP1 was used to inhibit the analogue-sensitive PKA allele (TPKas) in PKAi cells and was added at time point 0 h. Scale bars: 10 μm. **(B)** Quantification of cytoplasmic dry mass density from (A), where each cell was segmented into the cytoplasm and vacuole, and the vacuole was excluded in the dry mass density measurement. Mean ± SD of WT N = 274, PKAi N = 319, PKAi *msn2/4Δ* N = 272 cells. One representative dataset of two biological replicates per condition is shown. **(C)** Buoyant density measured by microcapillary density gradient centrifugation. Mean ± SD of N = 3 biological replicates. **(D, E, F)** Single-particle tracking of 40-nm GEMs was performed after 3 hours of PKA inhibition to determine mean-square displacement (MSD) curves and derive the diffusion coefficient, the percentage of unconfined trajectories and the radius of confinement (N = 3 biological replicates, mean ± SD). Statistics: One-Way ANOVA with Tukey’s multiple comparisons test (^ns^p > 0.05, *p ≤ 0.05, ***p ≤ 0.001, ****p ≤ 0.0001).

To measure the impact of these massive changes in mass density on the movement of macromolecules, we performed single-particle tracking using <u>g</u>enetically <u>e</u>ncoded <u>m</u>ultimeric nanoparticles (GEMs) ^8^. GEM trajectories were used to calculate time–ensemble-averaged mean-squared-displacement (TE-MSD) curves, from which the short-lag effective diffusion coefficient was obtained for each condition. Three hours of PKA inhibition led to a drastic drop in the diffusion coefficient of approximately 60% (**Figure 1D and S1D**). This is consistent with the notion that increased biomass density slows down particle mobility.

To get further mechanistic insight into how increased density might affect particle movement, we grouped our single particle trajectories based on the anomalous diffusion exponent *α* into groups that we termed confined (*α* < 0.8) or unconfined (*α* ≥ 0.8) as described previously ^18^. This classification revealed that inhibition of PKA reduced the percentage of unconfined trajectories by around 20% (**Figure 1E**) and the radius of confinement by close to 60% (**Figure 1F**). While particles classified as confined generally diffuse more slowly, the drop in diffusion upon PKA inhibition is similar for confined and unconfined trajectories (**Figures S1E and S1F**). This suggests that PKA inhibition increases crowding throughout the cytoplasm and not just in distinct pockets as shown in glucose depletion ^18^. Therefore, PKA-inhibited cells exhibit a strong, general decrease in diffusion and increase in confinement.

Inactivation of PKA leads to the activation of the transcription factors Msn2/4 and the subsequent differential regulation of hundreds of stress responsive genes ^4,5^. To determine whether this transcriptional response is required for the increase in cell density, we deleted *MSN2/4* **(Figure S1G)**. Indeed, the absence of *MSN2/4* prevented the density increase upon PKA inhibition (**Figures 1A, 1B and 1C**) and restored the GEM diffusion to wild-type (WT) levels (**Figures 1D, 1E, 1F, S1E and S1F**).

In contrast, hyperactivation of Msn2 through overexpression of a constitutively active *Msn2(S6A)* allele ^36^, increased cell density, although this effect is less pronounced than the density increase observed after PKA inhibition (**Figure S1H**). Activating the ESR through heat shock also led to an increase in buoyant density (**Figure S1I**). Together, these results show that PKA regulates density and diffusion via Msn2/4 and the ESR.

### PKA inhibition causes massive carbohydrate accumulation

Next, we wanted to determine the molecular makeup of the dense and crowded cytoplasm of PKA-inhibited cells. Since PKA inhibition did not decrease the cell size (**Figure S2A**), the rise in density is likely caused by an overproduction of macromolecular content. To test the role of protein synthesis, we inhibited translation elongation with cycloheximide (CHX). This treatment prevented the dry mass density increase (**Figures 2A and 2B**) and reduced the buoyant density increase in PKA-inhibited cells (**Figure 2C**). CHX treatment alone caused a mild increase in buoyant density (**Figure S2B**). The density increase caused by PKA inhibition is therefore dependent on the translation of proteins; either through downstream effects caused by stress response activation or because protein accumulation directly contributes to the density increase. To test the contribution of the latter, we measured bulk protein content normalized to the cell volume. PKA-inhibited cells showed a slight increase in protein concentration from 71 to 86 mg/mL compared to WT cells (**Figure 2D**). However, this increase did not reach significance and cannot account for the 90 mg/mL dry mass density increase measured after three hours of PKA inhibition **(Figure 2D)**. A general overproduction of proteins does therefore not explain the increased cell density observed after PKA inhibition.

**Figure 2.**
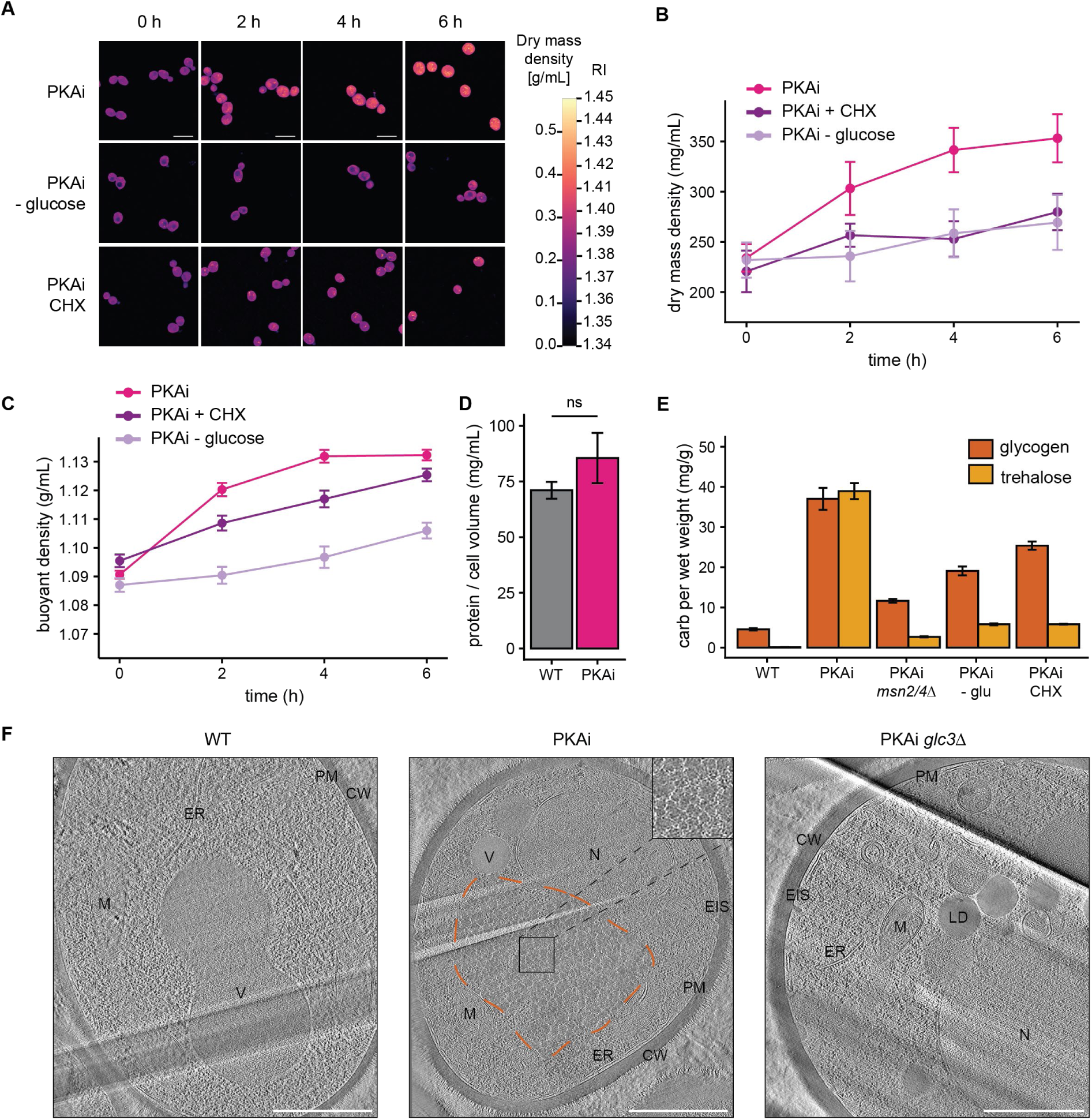
Cell density increase upon PKA inhibition coincides with carbohydrate accumulation. **(A)** Refractive index holotomography and conversion of refractive index into dry mass density of PKA inhibited cells, PKA inhibited cells grown without glucose in the medium or PKA inhibited with added cycloheximide (CHX). Glucose withdrawal and cycloheximide addition were performed at time point 0 together with the addition of PKA-inhibitor 1NM-PP1. Scale bar: 10 μm. **(B)** Quantification of (A). Mean ± SD of PKAi N = 558, PKAi + CHX N = 155, PKAi-glucose N = 462 cells. One representative dataset of two biological replicates per condition is shown. **(C)** Buoyant density measured by micro capillary density gradient centrifugation. Mean ± SD of N = 3 biological replicates. **(D)** Whole-cell protein content was measured by Bradford assay and normalized to cell volume measured by coulter counter in WT and PKAi cells 3 hours after addition of 1NM-PP1. Mean ± SD of N = 3 biological replicates. Statistics: Welch’s t-test (^ns^p > 0.05). **(E)** Trehalose and glycogen content was measured and normalized to wet weight of cells. Mean ± SD of N = 3 biological replicates. **(F)** Representative projections of cryo-Plamsa Focus Ion Beam (cryoPFIB)-milled untreated WT, PKA-inhibited and PKA-inhibited *glc3Δ* yeast cells after 3 h of PKA inhibition. N: nucleus; V: vacuole; M: mitochondrion; ER: endoplasmic reticulum; CW: cell wall; PM: plasma membrane; LD: lipid droplet; EIS: eisosome. The dashed orange line indicates an area of dense glycogen, magnified in the inset in the top right corner. Scale bars: 1 μm.

Since PKA is an important regulator of glucose metabolism, we next tested the contribution of glucose to the density increase of PKA-inhibited cells. Removal of glucose concurrent with PKA inhibition largely prevented the density increase (**Figures 2A, 2B and 2C**), demonstrating that the density increase requires glucose either as an energy source or as a building block. PKA inhibition and the ESR are known induce the production of the storage carbohydrates glycogen and trehalose ^37^. We therefore measured the weight fraction of glycogen and trehalose to determine whether their accumulation contributes to the density increase upon PKA inhibition. We measured a stark increase in both glycogen and trehalose levels in PKA-inhibited cells (**Figure 2E**). After PKA inhibition for three hours, glycogen and trehalose together constituted 76 ± 5 mg per gram of wet weight, accounting for most of the observed dry mass density increase of around 90 mg/mL at this time point (**Figure 1B**). Importantly, accumulation of these carbohydrates was greatly reduced in all the conditions that prevented the density increase: deletion of *MSN2/4*, glucose depletion or CHX treatment (**Figure 2E**). We conclude that glycogen and trehalose play a central role in the density increase upon PKA inhibition.

To gain further insight into how carbohydrate accumulation might influence the organization of the cytoplasm, we performed label-free *in situ* cryo-electron tomography (cryo-ET) on cells in a in a near native, hydrated state. PKA-inhibited cells displayed prominent clusters of dense granules in the center of the cytoplasm (**Figure 2F**) that are reminiscent of previously described glycogen rosettes ^38,39^. These structures were absent in cryo-electron tomograms of PKA-inhibited cells in which the gene encoding the glycogen branching enzyme Glc3 was deleted and glycogen synthesis was abrogated (**Figure 2F**) ^40,41^. Consistent with this, fluorescently tagged glycogen synthase Gsy2-mCherry localized to a similar central area in PKA-inhibited cells as the glycogen rosettes (**Figure S2C**). In contrast, PKA-inhibited *glc3Δ* cells showed a single Gsy2-mCherry focus (**Figure S2C**), which is characteristic of low glycogen levels ^42^. This strongly suggests that the rosettes observed in cryo-ET are glycogen and that it accumulates in the cytoplasm in vast quantities.

The cryo-electron tomograms also revealed that the organization of the cell surface and the cell interior changed significantly upon PKA inhibition. The cell wall was thicker in PKA-inhibited cells and the plasma membrane formed invaginations reminiscent of eisosomes ^43^. Additionally, in PKA-inhibited cells, the nuclei appeared deformed, the mitochondria fragmented and the vacuoles multilobulated and collapsed (**Figure 2F**). The latter two observations were confirmed by fluorescence microscopy with Tom70-mCherry to visualize the mitochondria and Vph1-GFP to visualize the vacuole (**Figures S2D and S2E)**. These deformations were observed proximal and distal to the glycogen granules, supporting our finding that the cytoplasm is more crowded everywhere, not just in distinct pockets.

Together, our results show that glycogen and trehalose accumulate in vast quantities in the cytoplasm of PKA-inhibited cells, impacting intracellular organization.

### Glycogen crowds the cytoplasm and contributes to density increase

To examine whether these carbohydrates contribute to the altered cytoplasmic properties in cells after PKA inhibition, we mutated genes required for their synthesis. To disrupt glycogen synthesis, we deleted *GLC3* as described above and in addition, also tested the deletion of the glycogenin glucosyltransferase encoding genes *GLG1/2* ^44^. These deletions did not affect the baseline glycogen levels or buoyant density (**Figures S3A and S3B**), but all led to a similarly diminished accumulation of glycogen upon PKA inhibition (**Figures 3A and S3A**). PKA inhibition in *glc3Δ* mutant cells still led to a density increase, but it was less strong than PKA inhibition in a WT background (**Figures 3B, 3C, 3D, and Table 1**). Therefore, glycogen contributes to the density increase upon PKA inhibition, but it is not the sole density-modulating component.

**Figure 3.**
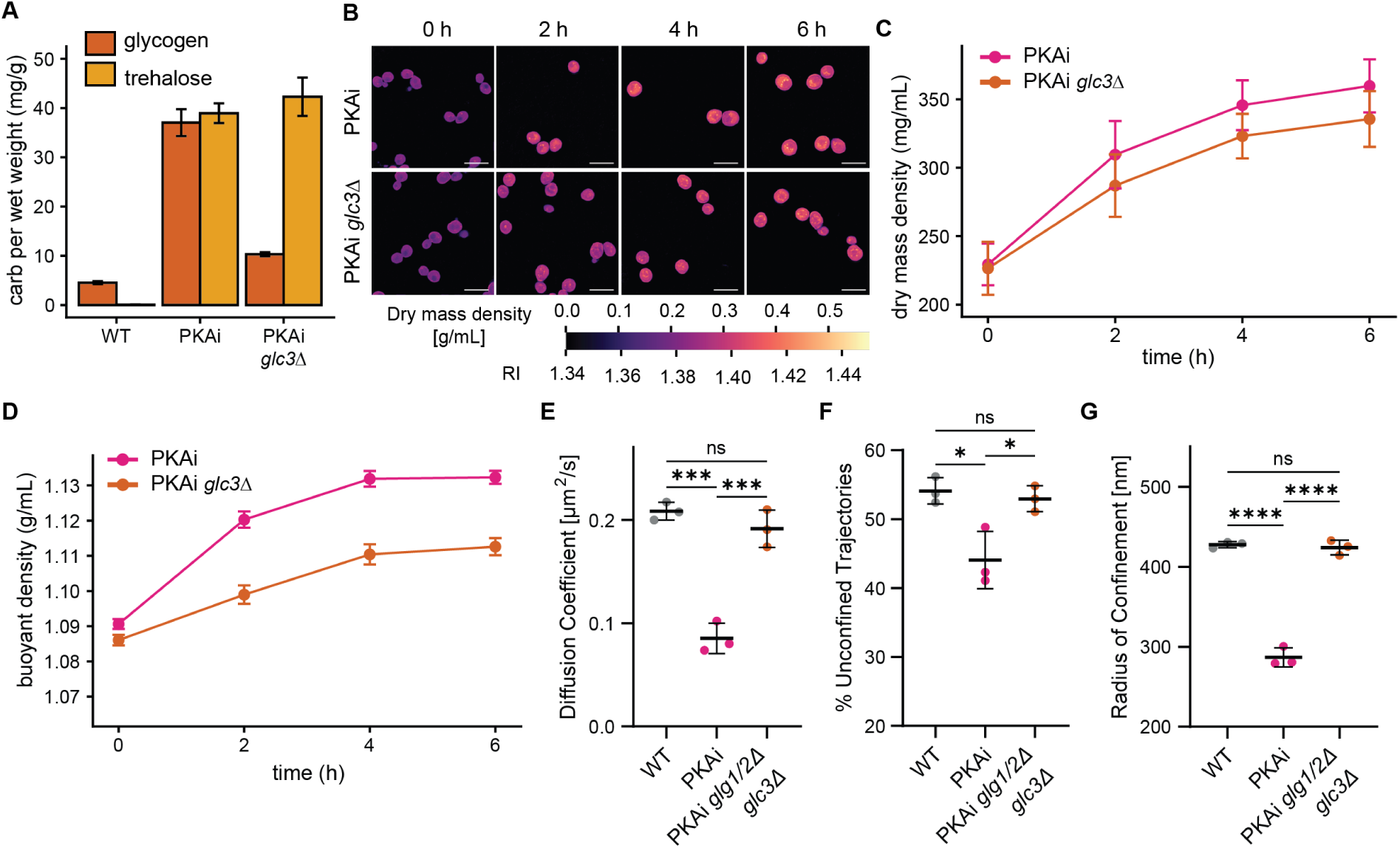
Glycogen reduces diffusion and contributes to cell density increase. **(A)** Trehalose and glycogen contents were measured and normalized to the wet weight of cells of the indicated genotypes 3 h after 1NM-PP1 addition. N = 3 biological replicates. **(B)** Refractive index holotomography was performed for cells of the indicated genotypes. 1NM-PP1 was added to inhibit PKA at time point 0 h. Scale bar: 10 μm. **(C)** Quantification of (B). Mean ± SD of PKAi N = 207, PKAi *glc3Δ* N = 302 cells. **(D)** Buoyant density of cells of the indicated genotype was measured by micro capillary density gradient centrifugation. PKA was inhibited by addition of 1NM-PP1 at time point 0 h. Mean ± SD of N = 3 biological replicates. **(E, F, G)** After 3 hours of PKA inhibition, GEM-expressing cells were imaged and the diffusion coefficient, the percentage of unconfined trajectories and the radius of confinement were determined (N = 3 biological replicates, mean ± SD). Statistics: One-Way ANOVA with Tukey’s multiple comparisons test (^ns^p > 0.05, *p ≤ 0.05, ***p ≤ 0.001, ****p ≤ 0.0001.

**Table 1:**
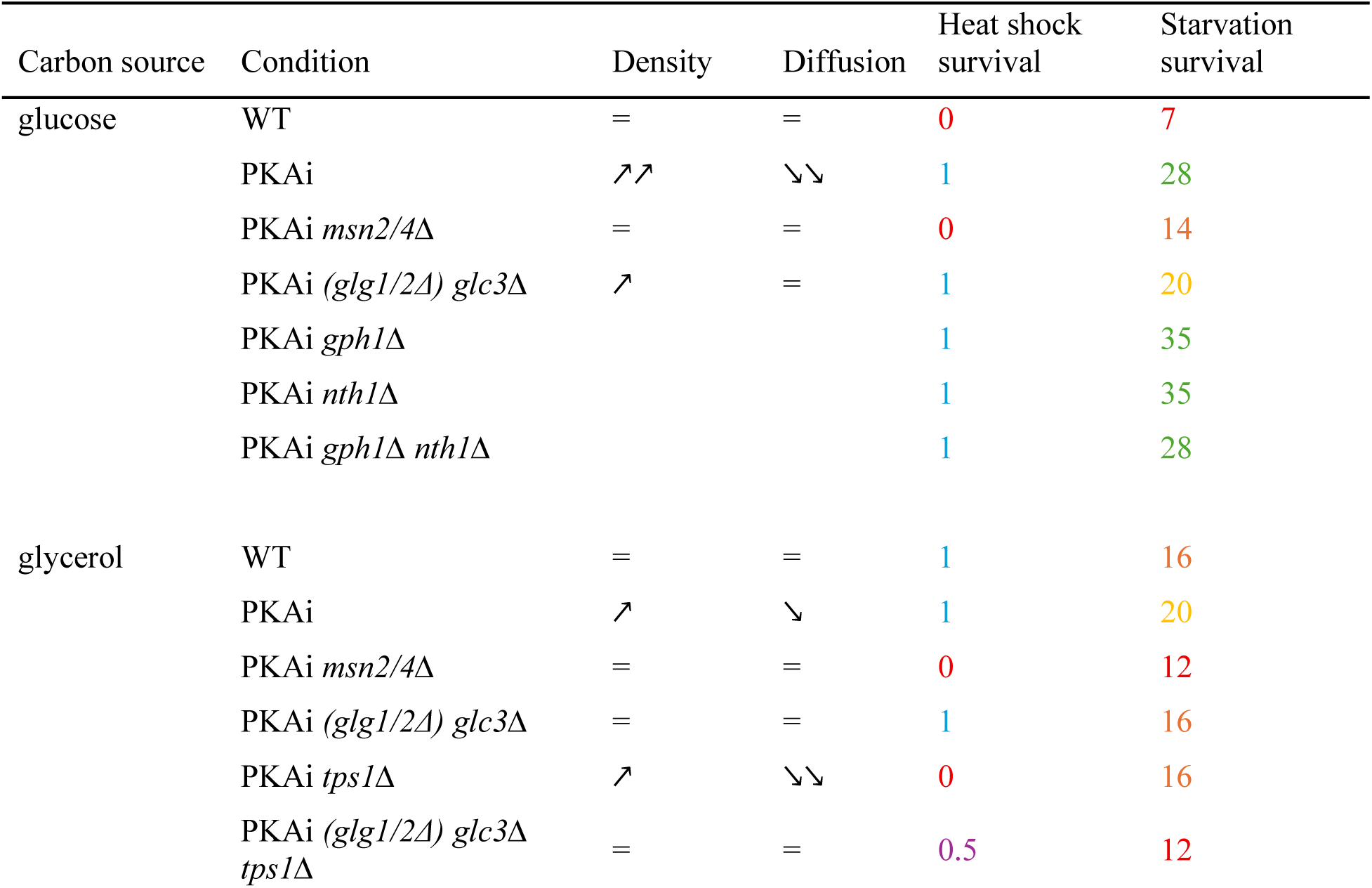
Overview of density, diffusion, heat shock survival and starvation survival in different conditions. Icons indicating behavior of density and diffusion, upon addition of the analogue 1NM-PP1: = unchanged, ↗ increase, ↘ decrease, ↘↘ stronger decrease. Survival of the tested heat shock conditions indicated by 1 for good survival and 0 for no or bad survival. Survival of the tested starvation conditions indicated by number of days after which significant decrease in regrowth was observed, color coded from green for long survival to red for short survival in weekly increments.

Interestingly, while disruption of glycogen synthesis only partially prevented the density increase upon PKA inhibition, it restored GEM diffusion and confinement to WT levels (**Figures 3E, 3F, 3G, S3C, S3D, S3E, and Table 1**). These results show that glycogen is the main contributing factor behind the slow and confined diffusion in PKA-inhibited cells. In addition, these measurements highlight that dry mass density, and diffusion properties are distinct features that need to be measured independently.

### Trehalose fluidizes the cytoplasm in crowded conditions

We next analyzed the contribution of trehalose to density and diffusion. Disruption of the trehalose synthesis pathway through deletion of the gene encoding trehalose-6-phosphate synthase *TPS1* causes growth defects in medium with fermentable carbon sources ^45^, and therefore all experiments with a *TPS1* deletion mutant were conducted in medium containing glycerol instead of glucose. In these conditions, logarithmically growing cells contained higher baseline levels of trehalose and glycogen (**Figures 4A and S4A**), likely due to reduced PKA activity under respiratory growth, as reported previously ^46^. PKA inhibition still led to an increase in both carbohydrates (**Figure 4A**), although the concentrations of glycogen and trehalose remained lower than in cells grown in glucose (**Figure 3A**). Deletion of *TPS1* fully abrogated trehalose production upon PKA inhibition, while *GLC3* deletion resulted in a substantial but only partial reduction of glycogen levels (**Figure 4A**). However, we consider it unlikely that the residual “glycogen” measured in the *GLC3* deletion mutant is actually glycogen, since the cryo-electron tomograms of this mutant showed no visible glycogen granules (**Figure 2F**) and even the mutant with the most severe impairment of glycogen synthesis (PKAi *glc3Δ glg1/2Δ* in **Figure S3A**) still exhibited residual “glycogen” levels. Instead, this analysis likely measures other glucans, such as the beta glucans of the thickened cell wall, that can be hydrolyzed in the acidic conditions of the glycogen extraction protocol ^47^.

**Figure 4.**
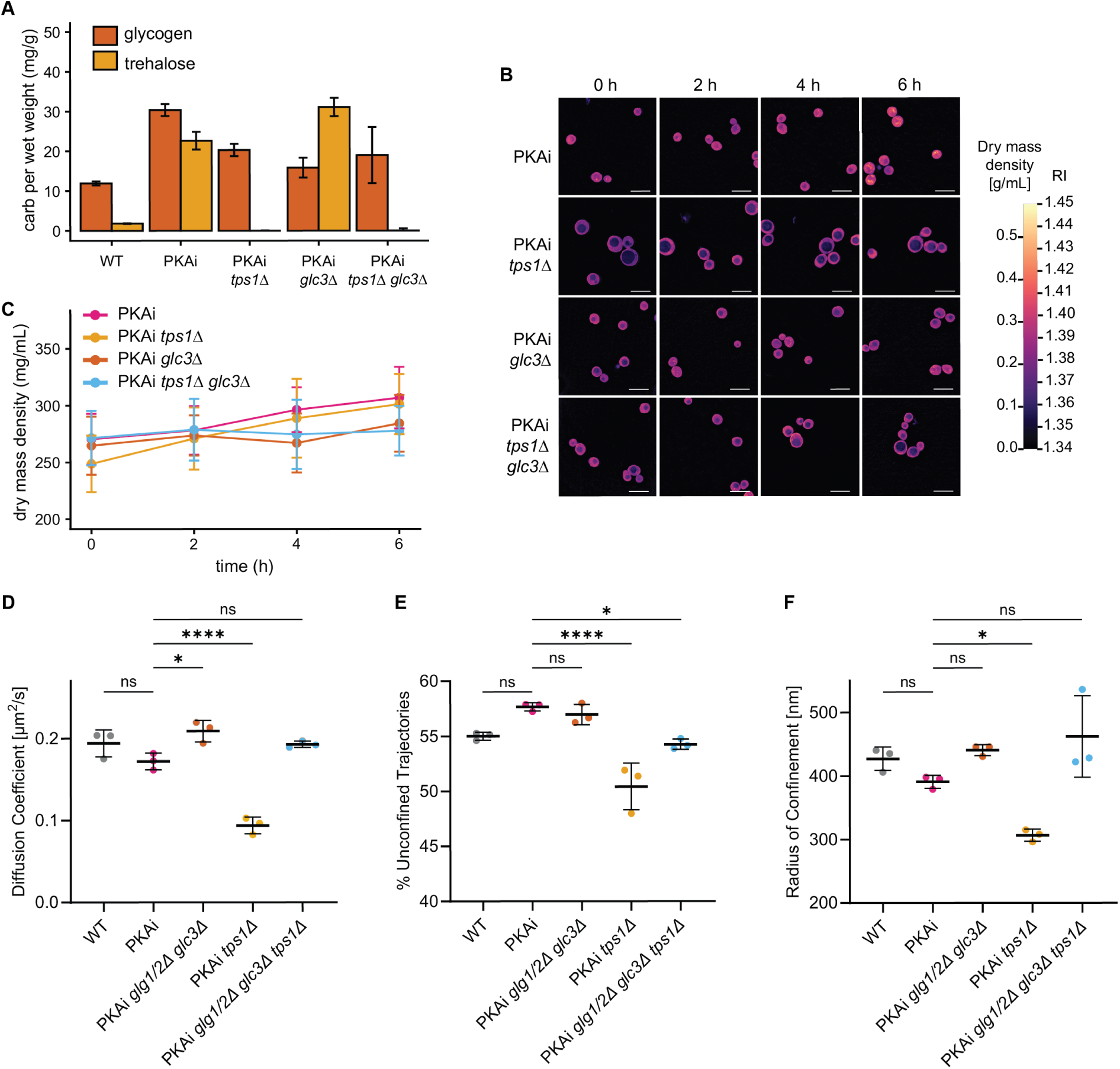
Trehalose fluidizes the cytoplasm upon PKA inhibition. **(A)** Trehalose and glycogen contents were measured and normalized to wet weight of cells of the indicated genotype 3 h after addition of 1NM-PP1 to inhibit PKA. Mean ± SD of N = 3 biological replicates. **(B)** Refractive index holotomography was performed for cells of the indicated genotypes. 1NM-PP1 was added to inhibit PKA at time point 0 h. Scale bar: 10 μm. **(C)** Quantification of (B). Mean ± SD of PKAi N = 223, PKAi *tps1Δ* N = 177, PKAi *glc3Δ* N = 204, PKAi *tps1Δ glc3Δ* N = 164 cells. One representative dataset of two biological replicates per condition is shown. **(D, E, F)** After 3 hours of PKA inhibition, GEM-expressing cells were imaged and the diffusion coefficient, the percentage of unconfined trajectories and the radius of confinement were determined (N = 3 biological replicates, mean ± SD). Statistics: One-Way ANOVA with Tukey’s multiple comparisons test (^ns^p > 0.05, *p ≤ 0.05, **p ≤ 0.01, ***p ≤ 0.001, ****p ≤ 0.0001), only discussed statistics shown.

Consistent with the carbohydrate measurements in glycerol-based medium upon PKA inhibition, cells start at a higher cytoplasmic dry mass density, and the density increase upon PKA inhibition is proportionally smaller than in glucose-based medium (**Figures 4B, 4C, and Table 1**). Deletion of *TPS1* had no effect on dry mass density, while *GLC3* deletion reduced the dry mass density increase upon PKA inhibition in glycerol medium (**Figure 4C and Table 1**). The buoyant density measurements were confounded by the fact that deletion of *TPS1* caused vacuoles to become very large, preventing any meaningful conclusions from these measurements (**Figure 4B, S4B and S4C**). Together, these results show that glycogen also contributes to the dry mass density increase upon PKA inhibition in non-fermentable carbon sources, while trehalose has no effect.

As glycogen accumulation led to decreased GEM mobility upon PKA inhibition in glucose-based medium, we wanted to assess the role of both glycogen and trehalose in regulating cytoplasmic diffusion when cells are grown in glycerol. PKA inhibition caused a smaller drop in the diffusion coefficient that was not significantly different from WT and fully dependent on glycogen synthesis (**Figures 4D, S4D, and Table 1**). Confinement was similarly less affected by PKA inhibition in glycerol than it had been in glucose (**Figures 4E, 4F, S4E and S4F**). In contrast, PKA inhibition in *TPS1* mutants led to a further decrease in GEM mobility (**Figures 4D, 4E, 4F, S4D, S4E, S4F, and Table 1**). Strikingly, this effect could be fully reversed upon deletion of glycogen synthesis enzymes **(Figure 4D)**. These results therefore show that glycogen and trehalose play different and opposing roles in regulating cytoplasmic mobility. While glycogen increases crowding and slows down diffusion, trehalose promotes cytoplasm fluidity upon PKA inhibition. This raises the interesting possibility that glycogen production needs to be accompanied by trehalose synthesis to prevent potentially detrimental rearrangements in an overcrowded cytoplasm that could result in extremely low diffusion rates.

### Glycogen and trehalose confer stress resistance independent of their role as energy sources

Reduced PKA activity and activation of the ESR have been shown to confer resistance to multiple stresses and extend survival during starvation ^22,2,18^. We therefore sought to investigate how the altered cytoplasmic properties and the production of trehalose and glycogen upon PKA inhibition contribute to stress survival.

We first determined cell survival after extreme heat stress. PKA-inhibited cells displayed dramatically increased survival compared to WT cells when shifted from 30 °C to 55 °C for 7.5 minutes (**Figures 5A, 5B, S5A, S5B, and Table 1**). This survival advantage was lost upon perturbations that abrogated the density increase in PKA-inhibited cells such as CHX treatment, *MSN2/4* deletion or the absence of glucose. However, it was not dependent on either the synthesis or degradation of glycogen (**Figure 5A and Table 1**). When grown in glycerol medium, WT cells were more resilient against heat stress, similar to PKA-inhibited cells (**Figure 5B and Table 1**). Survival of PKA-inhibited cells in glycerol was markedly reduced by CHX treatment or *MSN2/4* deletion (**Figures 5B and Table 1**), which was also the case in WT cells treated with heat shock (**Figure S5B**). As in glucose medium, preventing glycogen synthesis did not affect survival (**Figures 5B, S5B, and Table 1**). In contrast, trehalose synthesis-deficient cells were more sensitive to heat shock, as has been previously shown ^29^ (**Figures 5B, S5B, and Table 1**). Additional abrogation of glycogen synthesis appears to mildly rescue the heat sensitivity of *TPS1* deletion in PKA-inhibited cells (**Figure 5B and Table 1**), suggesting that glycogen accumulation becomes toxic if cells are not able to produce trehalose at the same time. This effect is not observed in cells without PKA inhibition (**Figure S5B**), implying that this only becomes critical at very high glycogen concentrations.

**Figure 5.**
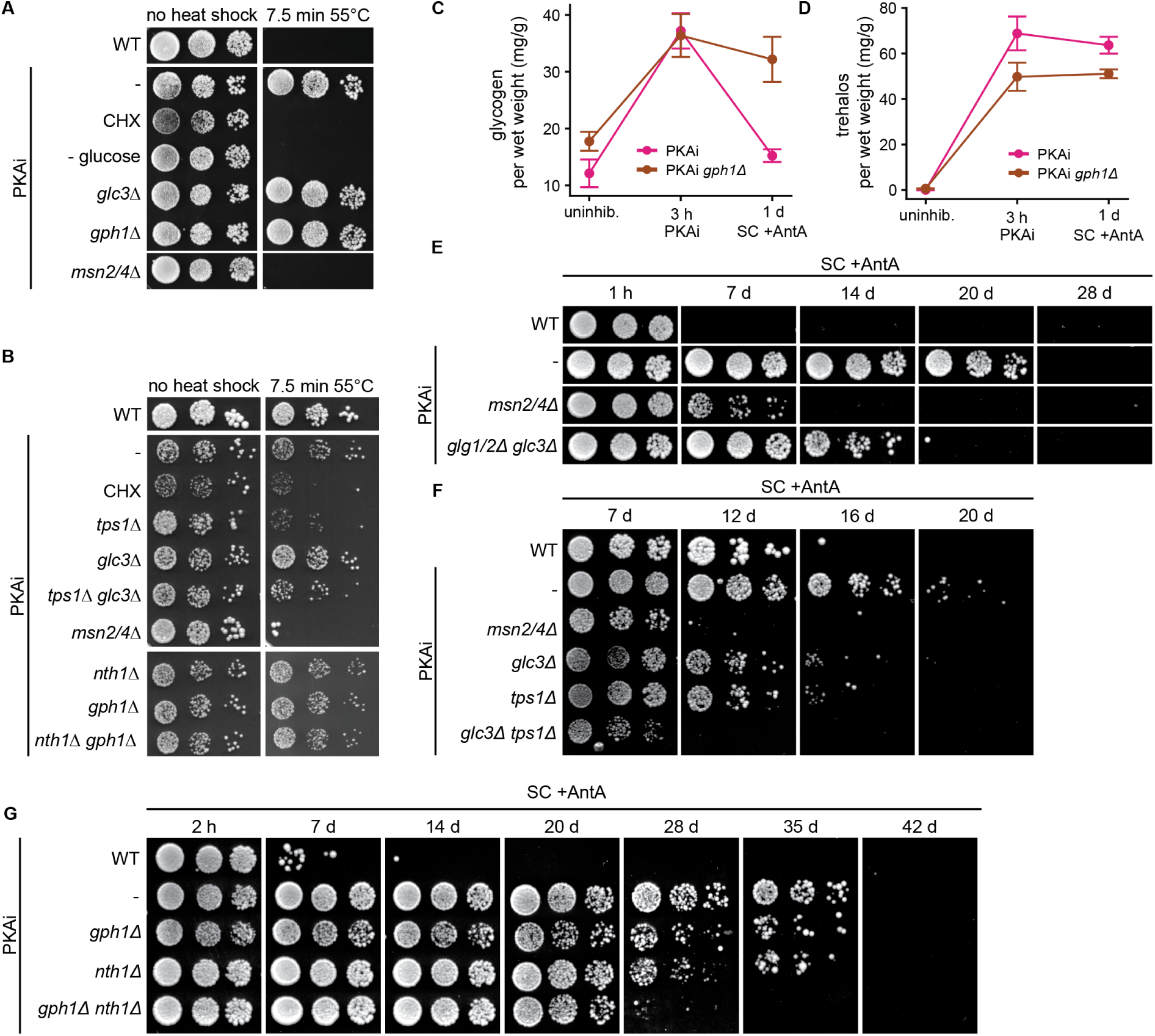
Glycogen and trehalose confer stress tolerance independent of their role as energy source. **(A, B)** PKA was inhibited in the indicated strains for 3 hours before cells were exposed to a heat shock from 30 °C to 55 °C for 7.5 minutes. Heat shock survival was assessed by plating 10-fold serial dilutions of cells before and after the heat shock. Cells were grown and plated and grown either on glucose-(A) or glycerol-(B) based plates. One dataset of two biological replicates is shown. **(C, D)** Glycogen (C) and trehalose (D) were measured in the indicated strains before treatment, after 3 hours of PKA inhibition and 1 day after removal of 1NM-PP1 and transfer of cells into starvation medium lacking glucose and containing Antimycin A (SC +AntA). Glycogen and trehalose contents were normalized to the wet weight of cells. N = 3 biological replicates. **(E, F, G)** Survival spotting of 5-fold dilution series of cells washed from glucose-(E, G) or glycerol-containing (F) medium into carbon-free medium containing Antimycin A (SC +AntA) for the indicated number of days. PKA was inhibited for 3 hours prior to transferring cells to SC +AntA. To assess viability qualitatively, cells were plated and grown either on glucose-(E,G) or glycerol-(F) based plates after indicated incubation times. One dataset of two biological replicates is shown.

To determine whether the critical role of storage carbohydrates for heat shock survival is to serve as an energy source, we deleted the enzymes needed for their degradation by mutating the neutral trehalase (*NTH1*) or glycogen phosphorylase (*GPH1*) (**Figures 5C and 5D**). Intriguingly, these mutations only had a moderate effect on heat shock survival in WT cells (**Figure S5B**) and no effect in PKA-inhibited cells (**Figures 5A and 5B**). We conclude that degradation of glycogen and trehalose is not essential for heat shock survival. Together these data show that accumulation of trehalose confers extreme heat stress resistance independent of its role as an energy source.

Yeast cells can survive very long periods without a carbon source, but cell survival declines rapidly upon abrupt glucose depletion when respiration is inhibited, and cells experience a complete lack of ATP production ^6,48^. Survival in these conditions is greatly improved when PKA is inhibited prior to respiration-deficient glucose starvation, and this effect strongly depends on Msn2/4 activity (**Figure 5E and Table 1**) ^18^. We therefore asked whether the increased glycogen- and/or trehalose levels contribute to this survival benefit. Survival spotting assays showed that deletion of the glycogen synthesis genes *GLG1/2* and *GLC3* significantly decreases survival of PKA-inhibited cells during energy depletion (**Figure 5E and Table 1**). When both glycogen and trehalose production are prevented in non-fermentable conditions, the entire growth benefit mediated by PKA inhibition is abolished (**Figure 5F**). The survival of these double mutants is now comparable to that of *MSN2/4* deletion mutants and even less than untreated WT cells, suggesting they too benefit from basal glycogen and trehalose levels present in growth on glycerol. Accumulation of glycogen and trehalose therefore greatly improves survival in energy-depleted conditions.

A simple explanation for this observation could be that storage carbohydrates serve as energy storage in nutrient-scarce conditions. However, the accumulated trehalose is not degraded in energy-depleted conditions and glycogen is degraded within one day of energy depletion, while the survival benefit lasts for weeks to come (**Figures 5C and 5D**). We therefore wondered whether storage carbohydrates might support survival independent of their role as energy source. Strikingly, preventing the degradation of glycogen or trehalose or both, by deleting *GPH1* and *NTH1* respectively, only had a minor effect on the starvation resistance provided by PKA inhibition (**Figure 5G**). These results demonstrate that glycogen and trehalose accumulation provide a survival benefit that does not depend on their role as an energy source. Rather, they suggest that the presence of glycogen and trehalose modulates the biophysical properties of the cytoplasm in a manner that is beneficial for surviving energy depletion.

## Discussion

In this work, we set out to determine how growth and stress signalling affect the material properties of the cell interior and whether these are important for the cellular adaptation to stress. We identify a critical role of PKA inhibition in regulating the biophysical properties of the cytoplasm by promoting the synthesis of glycogen and trehalose. While glycogen accumulation upon PKA inhibition leads to increased dry mass density and reduces the mobility of ribosome-sized particles, trehalose synthesis is needed to fluidize the crowded cytoplasm (**Figure 6**). Furthermore, we find that accumulation of these storage carbohydrates confers increased resistance to heat stress and to long term energy depletion. Importantly, heat resistance and prolonged survival of energy depletion do not depend on degradation of glycogen and trehalose. We therefore propose that glycogen and trehalose are not mere “storage-carbohydrates” as they have been viewed traditionally, but that they mainly promote stress and starvation survival by modulating cytoplasmic properties.

**Figure 6.**
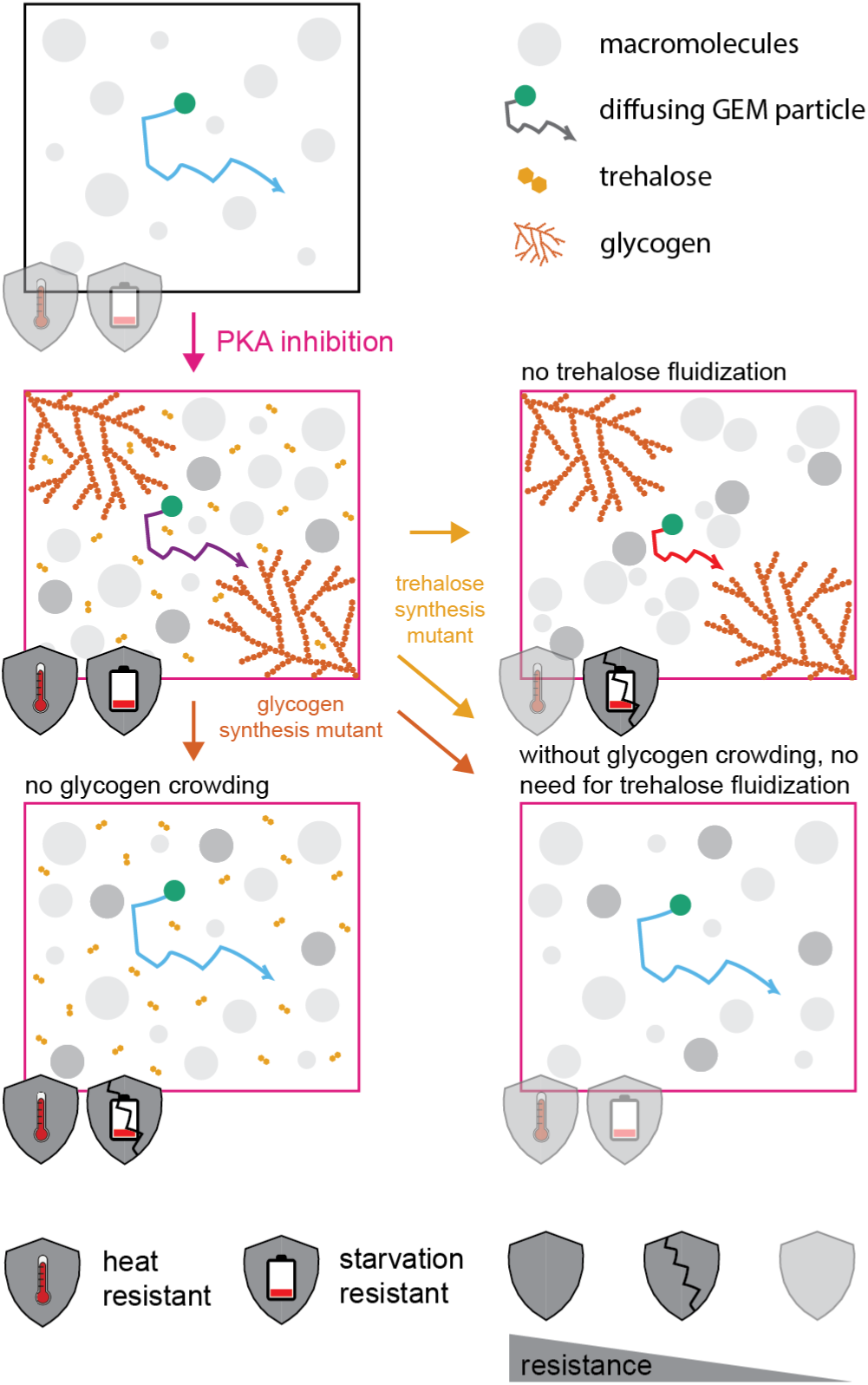
Inhibition of PKA increases stress resistance by altering the biophysical properties of the cytoplasm via accumulation of glycogen and trehalose. These carbohydrates grant resistance to heat and energy deprivation via distinct mechanisms. Glycogen partially drives the dry-mass density increase and strongly reduces the mobility of large (40 nm) GEM particles in the cytoplasm. The inability to produce glycogen severely compromises long-term survival in energy-depleted conditions (shield: starvation resistant). Conversely, trehalose does not alter dry-mass density but exerts a fluidizing effect that counters glycogen-induced particle immobilization. Without trehalose, heat resistance (shield: heat resistance) is entirely lost and energy depletion survival is diminished. Interestingly, double mutants lacking both carbohydrates bypass the need for trehalose’s fluidizing effect, yet they completely lose resistance to energy depletion while remaining partially heat-resistant.

Several methods to measure crowding-related phenomena have been developed in the last two decades ^49^. These measurements are often used interchangeably even though they rely on different physical measurements. Here we systematically compared three methods: The classical density gradient centrifugation to measure buoyant density, the more highly resolved refractive index holotomography to measure dry mass density and finally particle tracking with GEMs of 40 nm size to measure diffusion. In our experimentation, we identified conditions in which density and diffusion are decoupled. Specifically, PKA inhibition without glycogen synthesis increases density without reducing diffusion rates. Conversely, the lack of trehalose synthesis exacerbated the diffusion decrease upon PKA inhibition without further increasing density (**Figure 6**). The discrepancy between density and diffusion reflects the fact that density measurements include components across all size scales, while diffusion measurements are limited to the size scale affecting the tracer particle. The tracer used in this study is in the size range of ribosomes, which have been shown to greatly impact diffusion in the mesoscale depending on their abundance ^8,10,17^. Additionally, diffusion of tracer particles can be influenced by interactions with other molecules, by the size of these molecules and by how they are organized in space. Ultimately, our findings highlight that cytoplasmic density and macromolecular mobility are distinct biophysical properties that do not always correlate and require complementary evaluation.

The biophysical properties of the cytoplasm have previously been shown to respond sensitively to changes in the environment and stress. However, different environmental changes have different effects. Amino acid starvation and acute inhibition of TORC1 and translation have been shown to enhance diffusion in the cytoplasm. This change has been attributed to decreased crowding by reduced ribosome concentrations and polysome collapse ^8,50,16,17,51^. Glucose withdrawal on the other hand, which also leads to a halt of ribosome biogenesis and protein synthesis, leads to a dramatic reduction in macromolecular mobility ^6,7^. Here, we show that PKA, a major glucose-responsive growth regulator and modulator of stress signalling, can regulate the biophysical properties of the cell by regulating accumulation of glycogen and trehalose. We demonstrate that glycogen plays a role as a crowder. As such it is particularly interesting because it affects large parts of the mesoscale thanks to its variable chain length and branching structure resulting in particles ranging from 10 to 300 nm in diameter ^52^, comparable to sizes from proteasomes up to mitochondria. We find trehalose, though widely considered a viscogen that reduces diffusion and reaction rates ^30,11^, acts to fluidize the cytoplasm in conditions of high crowding. We propose that the cell regulates its biophysical properties through stress signalling by increasing crowding with glycogen and in parallel by keeping the cytoplasm fluid with trehalose.

PKA-inhibited cells share many properties with spores, such as elevated trehalose and glycogen levels, increased crowding, buoyant density, and overall tolerance to extreme stress ^53–55^. Our data suggests that the extreme tolerance to stress in spores is at least partially mediated by the altered cytoplasmic properties. However, stress resistance in spores has also been attributed to their massively thickened cell wall. Interestingly, we see a similar cell wall thickening in PKA-inhibited cells. This adaptation may be necessary to counteract increased internal pressure caused by the accumulation of osmotically active compounds such as trehalose. Without thickening of the cell wall, cells might expand instead of increasing density. Indeed, cell wall thickening has recently been shown to prevent cell wall stretching and allow crowding homeostasis when cells grow exceedingly large ^56^. Interestingly, increased cell wall thickness also contributes to heat resistance of spores ^53^. Whether the increased cell wall thickness itself prevents PKA-inhibited cells and spores from bursting or whether thicker cell walls are merely necessary to allow the cytoprotective accumulation of trehalose and glycogen remains to be determined.

Our data show that production of glycogen and trehalose confers stress resistance, but this effect does not require degradation of these carbohydrates, suggesting that they confer stress tolerance through another mechanism, potentially through the resulting altered cytoplasmic properties. Glycogen-mediated crowding could for example be beneficial by decreasing the rates of unserviceable and energy-consuming reactions. Because crowding differentially affects reactions based on the size of the interaction partners and the crowder, diffusion-limited reactions with larger interactors are disproportionately impacted ^57^. For example, translation decreases in overcrowded bacteria, yeast and human cells and was demonstrated to respond directly to overall crowding in *Xenopus* egg extracts ^12,58,14,59^. As translation involves megadalton-sized complexes, we speculate that increased crowding due to glycogen reduces rates of energy-costly translation ^60^, thereby preparing the cells for a quiescent state in which energy conservation is crucial.

Decreasing reaction rates can be beneficial in certain circumstances but excessive slowing down can also be detrimental. Our previous work has shown that a minimum level of fluidity is necessary to adjust the proteome during stress ^18^. Trehalose and glycogen production have been reported to contribute to the maintenance of invariant reaction rates upon heat stress and glucose deprivation ^11^. The fluidizing effect of trehalose we observe could therefore act to balance and mitigate the diffusion decrease imposed by glycogen crowding.

Beyond slowing reactions, crowding can also be cytoprotective by favoring phase separation and protein condensation through depletion-attraction forces ^61,8^. Cytoplasmic condensates (e.g. P-bodies, stress granules) have been found to sequester enzymes and substrates to modulate their activity in stress conditions ^62^. Glycogen acts as a crowder in our data and has also specifically been shown to contribute to cytoplasmic organization. In *Xenopus* egg extracts, glycogen-dependent demixing has been observed at increased crowding, where the extract demixes into a glycogen-rich and a glycogen-deplete phase, which sequesters mitochondria and ribosomes ^63^.

While crowding promotes functional condensation, it also affects aggregation dynamics of potentially harmful aggregates ^64–66^. The concomitant accumulation of trehalose might serve to protect proteins in glycogen-crowded conditions from aggregating by maintaining a hydration shell around them ^67,68^. Trehalose is also known to stabilize proteins, protecting them from denaturation, promoting correct folding and preserving enzyme activity ^32,69^. The need for trehalose as a fluidizing and stabilizing agent, specifically in highly crowded conditions, could explain the slight amelioration of heat resistance we see in glycogen and trehalose double mutants compared to trehalose mutants. It is possible that trehalose is particularly required to survive a heat shock as a fluidizing agent when glycogen accumulates in high levels and crowds the cell.

Together, our work highlights how cells can actively tune their internal material properties to withstand acute and chronic stress conditions by modulating carbohydrate metabolism. The roles of glycogen and trehalose are complementary and exceed their traditionally assigned roles as storage carbohydrates.

## Materials and Methods

### Strain generation

All *Saccharomyces cerevisiae* used in this study are derivatives of the W303 strain background. *S*trains were constructed either via transformation of linearized plasmids or PCR products with homology regions to the target site ^70^ or by crossing and microdissection. All strains, plasmids and primers used in this study are listed in Supplemental Tables S1–S3.

### Yeast cultivation

Growth media were prepared following standard protocols ^71^. For buoyant density measurements, refractive index tomography imaging, survival assays and protein, trehalose or glycogen extractions, cells were grown in yeast extract/peptone (YP) supplemented with adenine (0.055 mg/mL) and either 2% glucose (YPD) or 2% glycerol (YPG). For GEM tracking experiments, cells were grown in synthetic complete (SC) medium with 6.7g/l yeast nitrogen base (Difco), essential amino acids (Sigma, Merck), pH 5.0 and either 2% glucose or 3% glycerol and 2% ethanol. YPD plates consisted of 20 g/l bacto agar (BD Biosciences), 20 g/l bacto peptone (BD Biosciences), 10 g/l yeast extract (BD Biosciences), 2% w/v glucose (Sigma), and 80 mg/L adenine hemisulphate (Sigma). YPGlycerol plates contained the same ingredients, but 3% w/v glycerol instead of glucose. Unless stated otherwise, cells were grown at 30 °C overnight and diluted with fresh medium and grown into logarithmic phase (OD_600_ = 0.4) prior to any experiment.

### Drug treatments

Cycloheximide (CHX, Sigma) was dissolved in water and used at a final concentration of 20 μg/mL. 1-Naphthylmethyl PP1 (1NM-PP1, Sigma), dissolved in DMSO, was used at a final concentration of 5 μM and for the indicated time periods. Upon washing the cells into glucose-free medium, the 1NM-PP1 was washed out and not re-added. Antimycin A (Sigma) was dissolved in DMSO and used at a final concentration of 10 μM. For Msn2(S6A) overexpression, copper(II) sulfate pentahydrate (Sigma) was added to the culture at a final concentration of 10 μM.

### Stress survival assay

Resistance to extreme heat shock was determined as follows: Cultures were grown to logarithmic phase at 30 °C and treated as described. Prior to the heat shock, cell number was determined by measuring optical density (OD_600_) and cultures were aliquoted into 1.5 mL tubes. Cells were either left at room temperature or transferred to a water bath at 55° C for 7.5 minutes. After treatment, cells were diluted to OD_600_ = 0.2 and these cultures were then serially diluted either 10-fold or 5-fold three times. 5 µl or 3 µl of these dilutions were then plated onto YPD or YPGlycerol plates respectively, which were subsequently incubated at 30 °C for 2 or 4 days, respectively. The plates were imaged with a Vilber Fusion FX7 and processed in Fiji (ImageJ 1.54f) ^72^.

To determine cell survival during complete energy depletion, logarithmically growing cells were washed three times and released into synthetic complete medium lacking a carbon source and that was supplemented with Antimycin A to inhibit respiration (SC +AntA). To test how heat shock prior to energy depletion affects survival, logarithmically growing cells were incubated at 42 °C for 30 minutes, left to recover at 30 °C for 30 minutes and then washed into SC +AntA. After washing into starvation medium, cell number was determined by measuring OD_600_ and cells were incubated at 30 °C. At regular intervals after washing cells into starvation medium, cells were serially diluted (starting at an OD_600_ of 0.2) 5-fold three times and spotted onto YPD (4 µl) or YPGlycerol (2 µl) plates as described above.

### Microcapillary density gradient centrifugation

Density gradients were generated in microcapillaries (1.15 mm inner diameter, 75 mm length VITREX) by sequentially wicking drops of density medium containing increasing concentrations of colloidal silica into the capillary (LUDOX AM 30 wt. %, Sigma-Aldrich) diluted with 150 mM NaCl. Cells were isolated from logarithmically growing yeast cultures by centrifugation (2 min, 800 rcf). Media was removed carefully and the cells were resuspended in the lowest density droplet, which was wicked into the capillary first. Capillaries were then centrifuged in a microcapillary centrifuge for 2 min at 1500 rcf (Hettich HAEMATOKRIT 200). Images of capillaries were taken with back illumination (DÖRR LT-2020) using a Sony NEX-5N CCD camera with a SEL 1855 lens. The gray value profile along the capillary length was measured with Fiji (ImageJ 1.54f) ^72^. To extract absolute density values from the gray value distribution, we normalized the curve assuming a linear gradient between the lowest and the highest point on the capillary based on the density medium concentrations used. A standard curve was fitted to the gray value profile to determine the mean density of the population. Density beads from Cospheric LLC were used to visualize the range and separation of the method in Figure S1B.

### Live-cell microscopy

To determine dry-mass density via refractive index holotomography, cells were imaged on a Tomocube HT-2H QPI equipped with fluorescence imaging capabilities to acquire optical diffraction tomograms at 60X magnification with a water immersion objective, as described in ^35^. Cells were immobilized on Tomodishes (Tomocube) coated with 2 mg/mL Concanavalin A (Sigma) for 15 minutes and imaged in their respective growth medium.

Fluorescence images of the vacuole and glycogen synthase were performed on the Tomocube HT-2H using the same setup as described above. Vacuolar membrane protein Vph1 tagged with GFP (Vph1-GFP) was imaged at 30% intensity of the 470 nm LED for 200 ms. Glycogen synthase Gsy2 tagged with mCherry (Gsy2-mCherry) was imaged at 75% intensity of the 570 nm LED for 500 ms. Fluorescence images of mitochondria were taken on a Nikon Eclipse Ti-E at 100X with an oil immersion objective and 10% power of the 640 nm LED for 100 ms.

Imaging of 40 nm GEMs was performed as described in ^18^. In short, cells were imaged in 384 well glass bottom plates (Matrical) coated with concanavalin A (Sigma, C201) using a Nikon N-STORM equipped with a Hamamatsu Orca Flash 4.0 V3, a 100X SR Apochromat TIRF objective with 1.49 NA. Effective pixel size was 160 nm. Images were acquired in HILO mode at a frequency of 33 frames per second for 700 frames using NIS Elements Advanced software.

### Image analysis

Refractive index holotomograms were segmented to exclude the vacuole from the density measurement of the cytoplasm dry mass density. Vacuoles were visualized with vacuolar membrane protein Vph1 tagged with GFP (Vph1-GFP). A 2D fluorescence image was acquired in the center plane of the 3D refractive index holotomogram. The center slice of the holotomogram was matched to the fluorescence image and used to determine the dry mass density in the cytoplasm. Each cell outline was segmented based on the refractive index image, and the vacuole was segmented based on the vacuolar fluorescent signal using a custom pipeline in CellProfiler 4 ^73^. Pixels assigned to the vacuole were then excluded from the measurement of dry-mass density of the cytoplasm. Refractive index values were converted to dry-mass density using an RI increment of 0.1907 mL/g ^74^. From the same pipeline we also measured the 2D area occupied by the cell.

GEMs were localized and tracked using the TrackMate 7 plugin ^75^ of Fiji ^72^ (ImageJ 1.54f). The localization “Quality” threshold was adjusted to account for signal-to-noise variations across the experiments. Trajectory analysis was performed using custom MATLAB scripts as described in ^18^. Software is available in this repository https://github.com/PabloAu/Single-Molecule-Tracking-Analysis. The TE-MSDs were calculated from the two-dimensional (x,y) trajectories. The effective diffusion coefficient was estimated from a linear fit to the TE-MSD using the first 4 points. α coefficients were obtained from a power-law fitting of individual T-MSD using 20 points. Trajectories with α < 0.8 were classified as confined; the remaining trajectories were classified as unconfined. The confinement radius was obtained for the confined subset using a circle confined diffusion model ^76^ and should be interpreted as an effective in-plane confinement radius.

### Cryo-electron tomography

Yeast cell cultures were grown overnight in YPD medium on a shaker (180 rpm) at 30°C, diluted to OD600 = 0.05, recovered for 2 hours and then treated with 5 μM 1NM-PP1 for 3 hours. Samples were harvested at OD600 = ∼0.6 by spinning down for 2 min at 800 g at room temperature and concentrated to OD600 = ∼3 in 1ml of conditioned media. Yeast cells were vitrified by plunge freezing using an automatic Leica EM GP2 (Leica Microsystems) ^77,78^. 4 μL of cell suspension was applied onto negatively glow-discharged (twice, 25mA for 45 sec) copper EM grids (R2/2, 200 mesh, Quantifoil). Grids were blotted for 6 seconds from the backside at 8°C, 95% humidity, before plunging into the liquid ethane-propane mixture ^79^.

Vitrified cells on EM grids were processed using automated sequential FIB milling in an Arctis dual-beam cryo-Plasma Focused Ion Beam–Scanning Electron Microscope (PFIB-SEM; Thermo Fisher Scientific)^80,81^. Prior to milling, samples were sputter coated with platinum for 120 s at 12 kV and 70 nA, followed by platinum deposition using the gas injection system (GIS) for 150 s. A second platinum sputter coating step was then performed for 120 s under the same conditions. Relief cuts were milled at 30 kV and 1 nA using a depth correction of 200%. Automated lamella was set up to target a final lamella thickness of 150 nm and a width of 20 µm using a rectangular milling pattern. Rough milling was performed at 30 kV and 1 nA with 150% depth correction, followed by medium milling at 30 kV and 0.3 nA with 200% depth correction. Fine milling was carried out at 30 kV and 0.1 nA with 100% depth correction. Polishing was completed in two steps: first at 30 kV and 30 pA with 250% depth correction, and then at 30 kV and 10 pA with 200% depth correction. Cryo-PFIB-milled lamellae were imaged in a Titan Krios G4 transmission electron microscope (Thermo Fisher Scientific) operating at 300 kV and equipped with a Gatan GIF-quantum energy filter (20 eV slit width) and a K3 direct electron detector (Gatan). Tilt series were collected in SerialEM v4.2 using SPACEtomo v1.3.0 and PACE-TOMO v1.9.2 at a nominal magnification of 6,500x (13.71 Å/px) with a target defocus of –20 µm ^82^; https://github.com/eisfabian/SPACEtomo.git; ^83^. Data were acquired using a dose symmetric scheme from –51° to +69° with 3° increment, yielding a cumulative electron dose of approximately 150 e⁻/Å². Movie frames were aligned using *alignframes*, and tomograms were reconstructed manually in IMOD with a binning factor of 4, resulting in a final pixel size of 54.84 Å/px ^84^.

### Protein measurement

10 mL of cell culture were collected for protein extraction and 1 mL for volume measurement. For protein extraction, cells were pelleted for 2 minutes at 16’000 rcf, washed with 1 mL of cold 10 mM Tris-HCl pH 7.5 and frozen in liquid nitrogen. The pellet was resuspended in 100 μL Lysis buffer (50 mM Tris-HCl pH 7.5, 1 mM EDTA pH 8, 1x Halt Protease Inhibitor (Thermo Scientific), 50 mM DTT). An equal volume of acid-washed beads was added and the cells were broken in a Fast Prep cell disruptor (FastPrep-24 5G cell disruptor, speed 6.5 for 45 seconds, 3-5 cycles) until >90% of cells were broken as assessed under a microscope. Lysed cells were transferred to a new tube and diluted with 50 mM Tris-HCl pH 7.5 for quantification with Bradford (Bio-Rad Protein Assay). Cell volume was measured on a Multisizer 4e (Beckman) with a 100 μm aperture and Isotone II (Beckman) as a diluent. Total protein content was divided by total cell volume of the collected cells.

### Trehalose and glycogen extraction and measurement

We extracted trehalose and alkali- and acid-soluble glycogen with an altered protocol based on ^85^ and ^86^. 10 mL cell culture at OD_600_ = 0.5 were collected by centrifugation (2 min, 800 rcf), transferred to 21.5 mL screw top tubes and washed three times with water, which was carefully removed after the last wash by aspiration. The 1.5 mL collection tubes were weighed before and after cell collection to determine the wet weight of the pellet. Finally, pellets were flash frozen in liquid nitrogen.

Pellets were resuspended in 7.5% TCA (trichloroacetic acid, Carl Roth) and incubated on ice for 1 hour to lyse the cells. The released trehalose was collected by centrifuging for 30 seconds at 10’000 rcf, washing with 200 μL water and combining the supernatants. To extract alkali-soluble glycogen, pellets were resuspended in 125 μL 0.25M sodium carbonate, vortexed for 10 seconds, incubated at 85 °C on thermal shaker with 700 rpm for 4 hours. This procedure liberates alkali soluble glycogen into the supernatant, which can then be separated from the cells by centrifugation. Acid-soluble glycogen was extracted by resuspending the pellet in 125 μL 0.5 M perchloric acid, boiling the samples at 100 °C on thermal shaker with 700 rpm for 30 minutes and collecting the supernatant after spinning down the cell debris by centrifugation.

The glycogen samples were digested with amyloglucosidase overnight. For this, 75 μL of the sample were combined with 45 μL 1M acetic acid, 180 μL sodium acetate (200 mM, pH 5.2) and 1.2 μL amyloglucosidase (1000 U/mL, from Aspergillus niger, Roche) and incubated at 60 °C on a thermal shaker at 700 rpm for 16 hours. This treatment breaks down glycogen into its glucose subunits which can subsequently be measured using a commercially available kit (Trehalose Assay Kit, Megazyme, NEOGEN). The abundance of trehalose was measured by treating trehalose containing cell extract with trehalase to break it down into its glucose subunits, which was then measured with the same enzymatic kit. The obtained values were normalized to the wet weight of the collected cells and only acid-soluble glycogen was shown because alkali-soluble glycogen levels were low in all conditions.

## End notes

## Acknowledgments

The authors would like to thank the members of the Neurohr, Weis and Peter lab for discussions and comments on the manuscript. We would also like to acknowledge ScopeM for their support & assistance in this work.

This work was supported by grants from the Swiss National Science Foundation to K.W. (TMAG-3_209354 and CRSII5_193740), to G.N. (310030_212660) and to M.P. (SNF_236080).

## Author contributions

Marina Kunzi: conceptualization, data curation, formal analysis, investigation, visualization, methodology, writing – original draft, writing – review & editing. Lorena Kronig: conceptualization, data curation, formal analysis, investigation, visualization, methodology, writing – original draft, writing – review & editing. Pablo Aurelio Gómez-García: methodology, software. Martina Bonassera: methodology, resources, visualization. Matthias Peter: conceptualization, supervision, funding acquisition, project administration. Karsten Weis: conceptualization, supervision, writing – review & editing, resources, funding acquisition, project administration. Gabriel Neurohr: conceptualization, supervision, writing – review & editing, resources, funding acquisition, project administration.

## Supplementary information

**Supplementary Figure S1.**
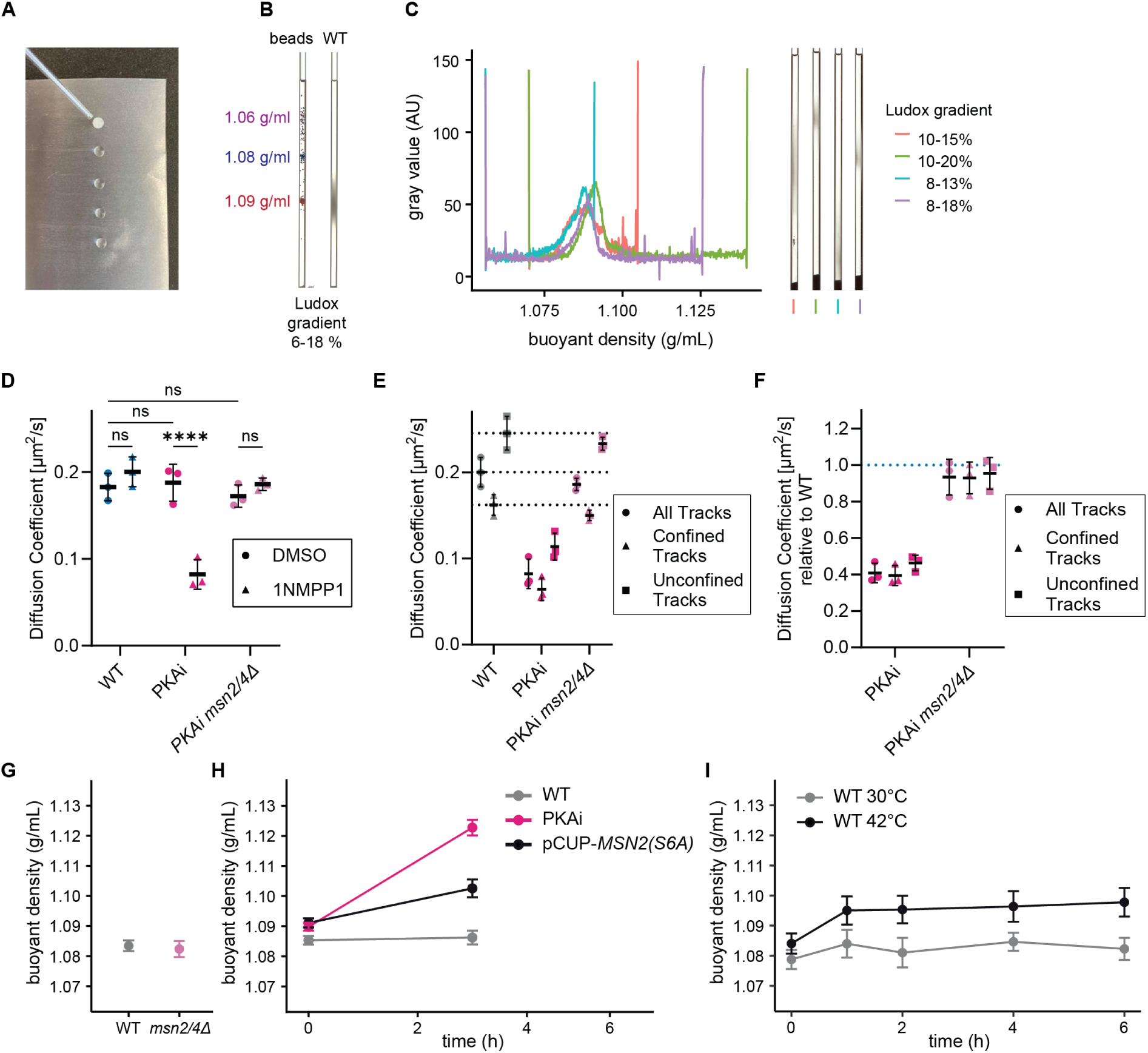
**(A)** Image of sample preparation for microcapillary density gradient centrifugation. The drops of density media dilutions, containing different concentrations of Ludox, are ordered from least dense (top) to densest (bottom) with the yeast sample resuspended in the least dense medium on parafilm. Drops are then sequentially aspirated into a microcapillary, starting with the least dense drop. **(B)** Demonstration of the range, spread and separation achieved with capillary density gradient centrifugation. Backlit images of capillaries containing the same density gradient. The left capillary contains density beads of different densities (1.06, 1.08, 1.09 g/mL), the right capillary contains WT yeast. **(C)** Gray value profiles of the capillaries shown on the right containing WT yeast separated on different density gradients. While flat gradients yield a wider cell distribution, the measured profiles almost collapse on each other when normalized to the gradient. These measurements show that buoyancy can be measured reproducibly across gradients with this simple method. **(D)** The indicated strains were treated for 3 hours with either DMSO or 1NM-PP1 and GEMs were imaged to determine the diffusion coefficient (N = 3 biological replicates, mean ± SD). Statistics: Two-Way ANOVA with Tukey’s multiple comparisons test (^ns^p > 0.05, ****p ≤ 0.0001). This control shows that treatment with 1NM-PP1 only affects GEM diffusion in an analogue-sensitive TPKas background (in PKAi strains) but not in WT cells. **(E)** The diffusion coefficient of all trajectories or specifically the confined (α < 0.8) or unconfined (α ≥ 0.8) trajectories were calculated from the MSD curves (N = 3 biological replicates, mean ± SD). The dotted lines indicate the WT values. **(F)** Diffusion coefficients of all, confined or unconfined trajectories were normalized to the respective WT values, showing that all subclasses of trajectories are similarly affected by inhibition of PKA. **(G)** Buoyant density measured by microcapillary density gradient centrifugation of the indicated strains. N = 3 biological replicates. WT data from Figure 1C is shown for comparison. **(H)** Buoyant density measured by microcapillary density gradient centrifugation of the indicated strains. Expression of the constitutively active *MSN2(S6A)* allele from a copper-inducible promoter was induced at time point 0. N = 3 biological replicates. (**I**) Buoyant densities of WT cells grown at 30 °C or shifted to 42 °C at the start of the experiment.

**Supplementary Figure S2.**
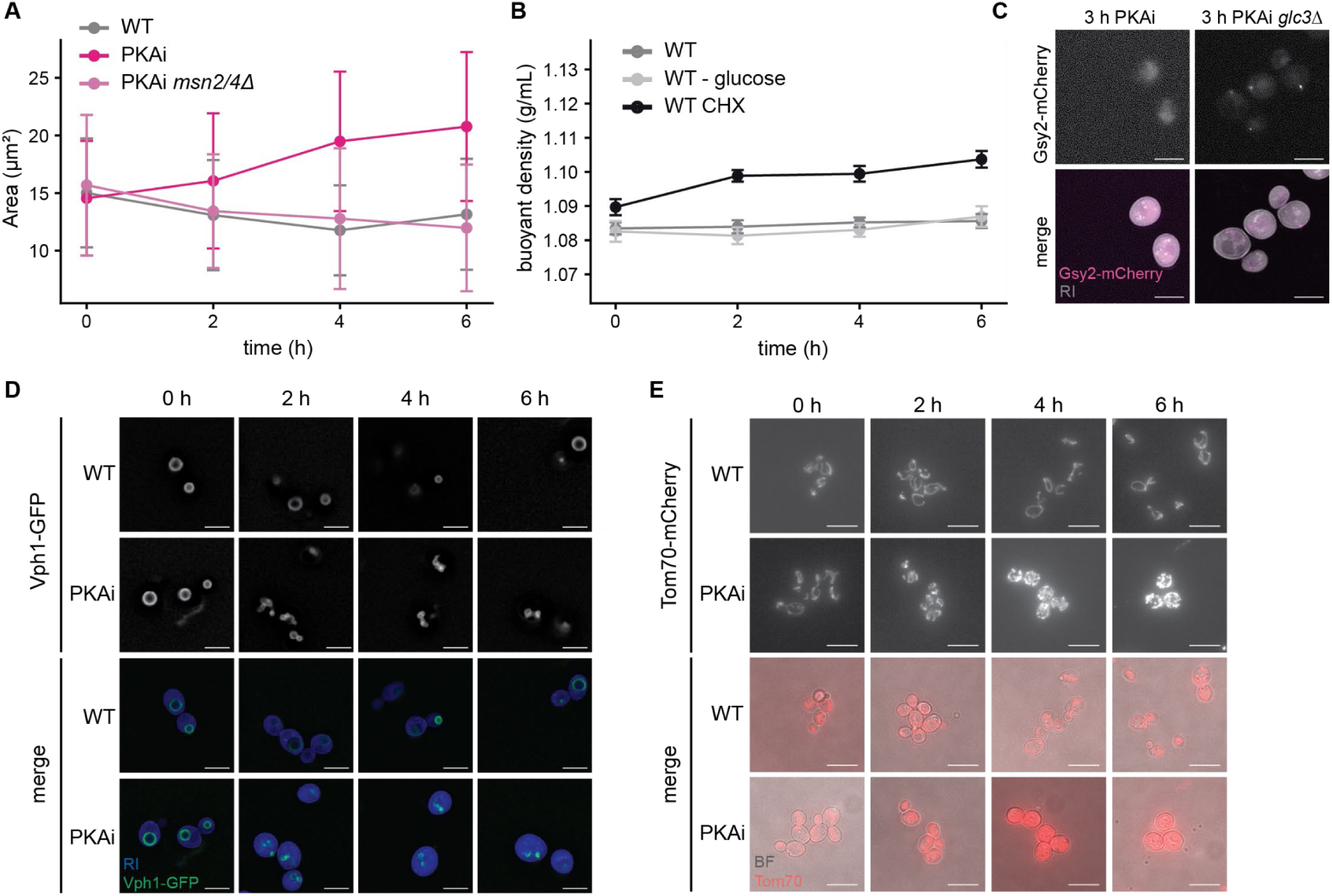
**(A)** Quantification of cell area measured in cells shown in Figure 1A. Average cell area measured from cross-sections of refractive index holotomography. **(B)** Buoyant density measured by micro capillary density gradient centrifugation. CHX was added at time point 0. N = 3 biological replicates. **(C)** PKA-inhibited cells expressing *Gsy2-mCherry* and *Vph1-GFP*, visualizing sites of glycogen synthesis and vacuolar membranes, respectively. Scale bar: 5 μm. (RI = refractive index) **(D)** Fluorescence microscopy images of cells expressing *Vph1-GFP* to visualize vacuolar membranes. Scale bar: 5 μm. (RI = Refractive Index). **(E)** Fluorescence microscopy images of cells expressing *Tom70-mCherry* to visualize mitochondria. Stationary phase sample of WT was taken. Scale bar: 10 μm. (BF = bright field).

**Supplementary Figure S3.**
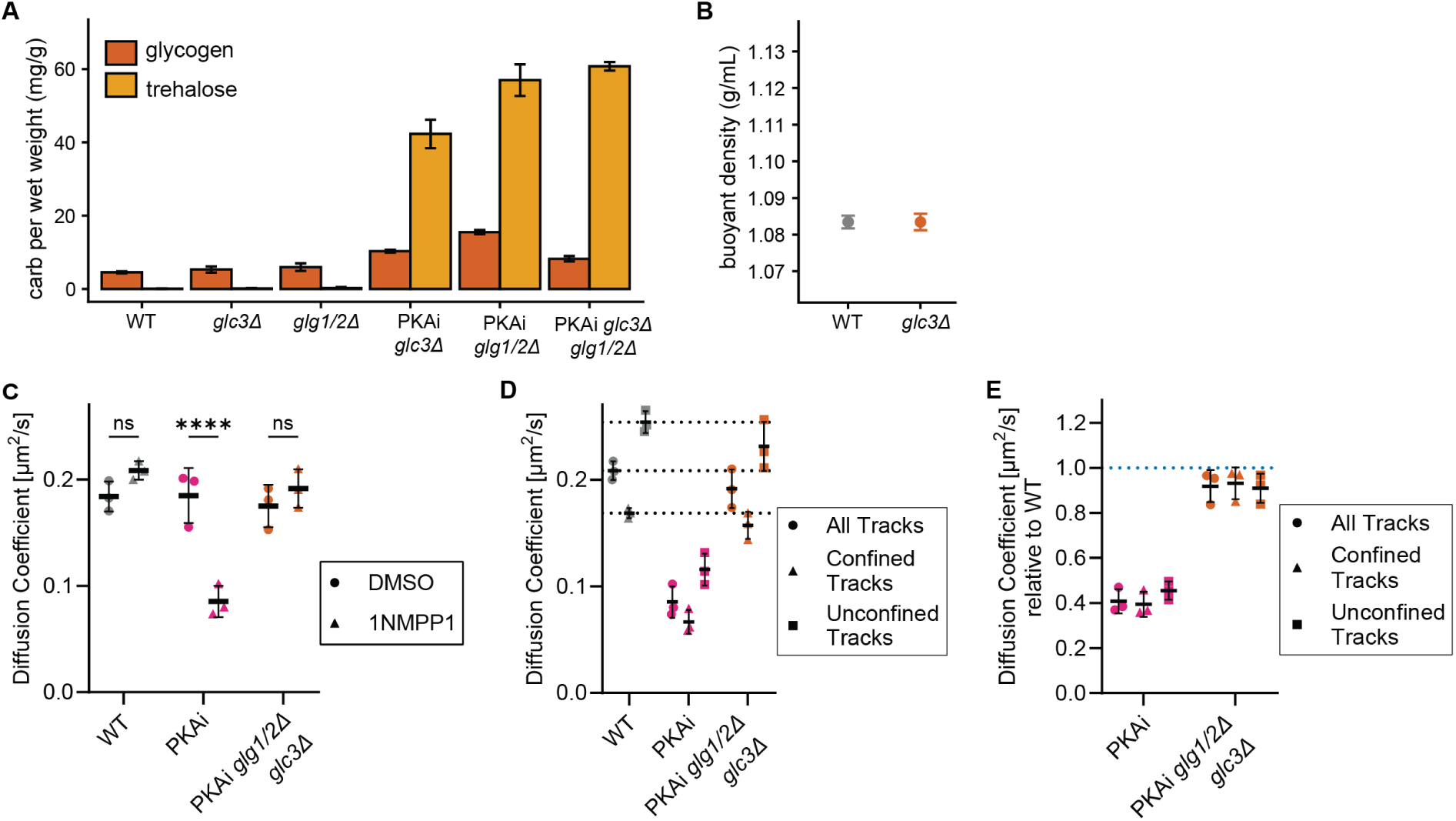
**(A)** Trehalose and glycogen contents in the indicated conditions were measured and normalized to wet weight of cells. N = 3 biological replicates. **(B)** Buoyant density measured by microcapillary density gradient centrifugation. N = 3 biological replicates. WT data from Figure 1C is shown for comparison**. (C)** After 3 hours of PKA inhibition, GEM-expressing cells were imaged and the diffusion coefficient was determined (N = 3 biological replicates, mean ± SD). Statistics: Two-Way ANOVA with Tukey’s multiple comparisons test (^ns^p > 0.05, ****p ≤ 0.0001). **(D)** The diffusion coefficients of the indicated trajectory subclasses were calculated from the MSD curves (N = 2 biological replicates, mean ± SD). The dotted lines indicate the WT values. **(E)** Diffusion coefficients of the indicated trajectory subclasses normalized to the WT values.

**Supplementary Figure S4.**
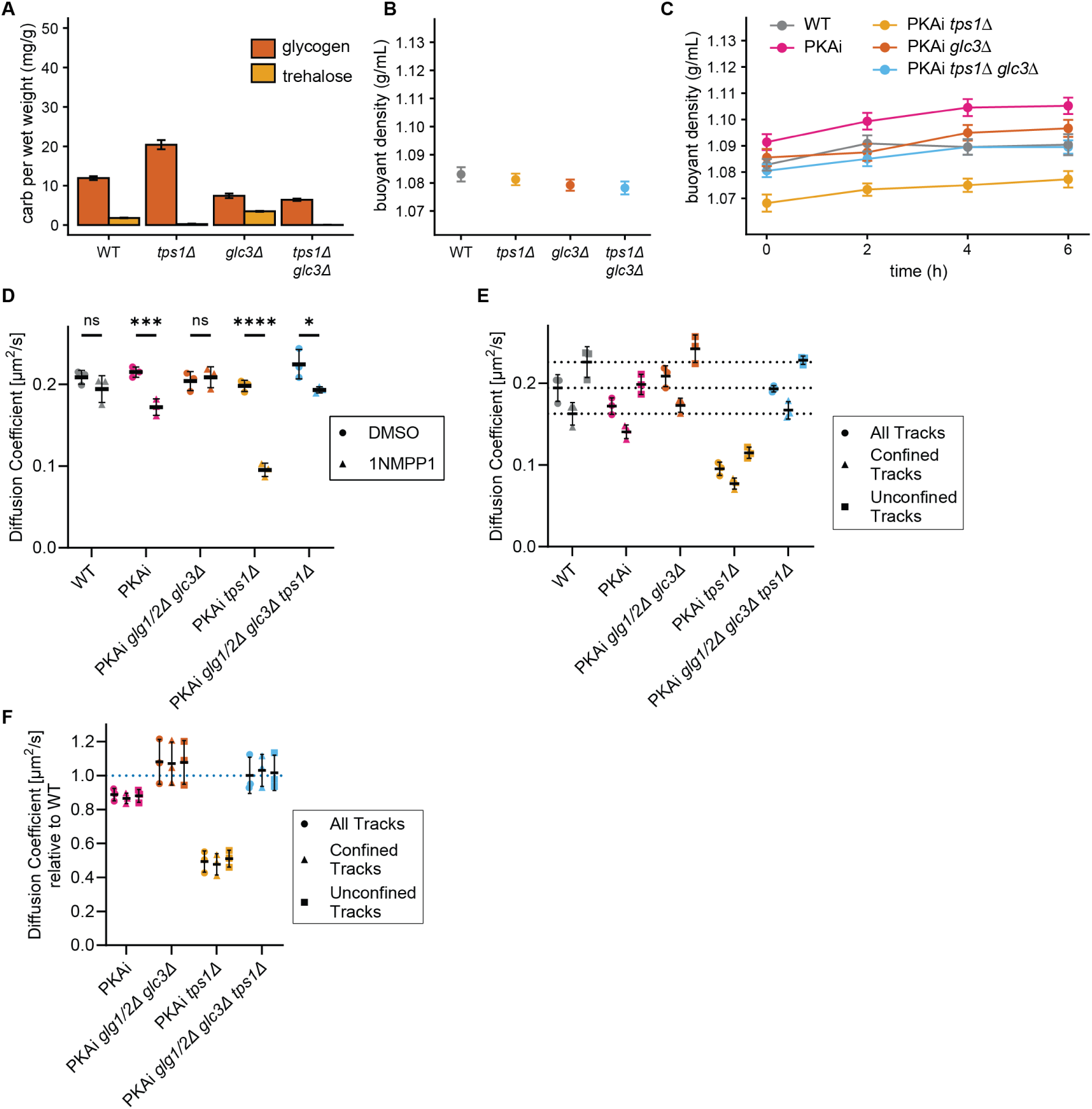
**(A)** Trehalose and glycogen contents in the indicated strains were measured and normalized to wet weight of cells. N = 3 biological replicates. Samples were extracted from cells grown in medium containing glycerol. **(B)** Buoyant density of the indicated strains measured by microcapillary density gradient centrifugation. N = 3 biological replicates. **(C)** Buoyant density measured by microcapillary density gradient centrifugation after inhibition of PKA using 1NM-PP1. N = 3 biological replicates. **(D)** Cells were grown in medium with glycerol and ethanol as carbon sources. After 3 hours of PKA inhibition, GEM-expressing cells were imaged and the diffusion coefficient was determined (N = 3 biological replicates, mean ± SD). Statistics: Two-Way ANOVA with Tukey’s multiple comparisons test (^ns^p > 0.05, *p ≤ 0.05, ***p ≤ 0.001, ***p ≤ 0.0001). **(E)** Cells were grown in medium with glycerol and ethanol as carbon sources. The diffusion coefficients of all trajectories and of the confined or unconfined trajectory subclasses were calculated from the MSD curves (N = 2 biological replicates, mean ± SD). The dotted lines indicate wild-type (WT) values. **(F)** Diffusion coefficients of indicated trajectory subclasses normalized to the WT values.

**Supplementary Figure S5.**
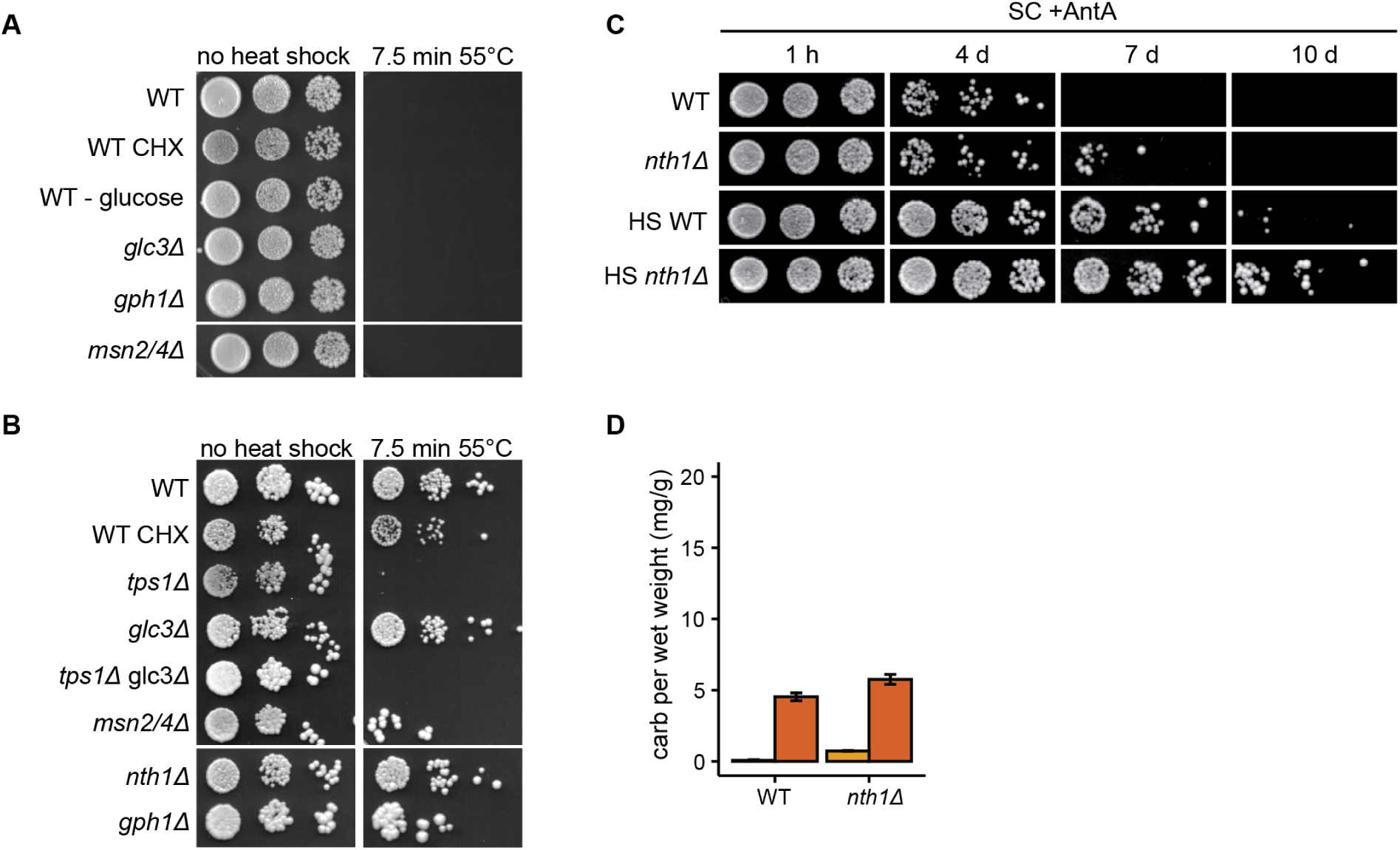
**(A, B)** Indicated strains were exposed to a heat shock from 30 °C to 55 °C for 7.5 minutes. Heat shock survival was assessed by plating 10-fold serial dilutions of cells before and after the heat shock. Cells were grown and plated either on glucose-(A) or glycerol-(B) based medium. **(C)** WT or *nth1Δ* cells were washed into starvation medium lacking glucose and containing Antimycin A (SC +AntA) with or without prior heat shock (HS, 42 °C for 30 min, 30 min recovery at 30 °C). To assess viability, 10-fold serial dilutions of cells were plated after the indicated incubation time and plates were imaged after 3 days at 30 °C. **(D)** Trehalose and glycogen contents normalized to wet weight of WT and *nth1Δ* mutant cells. N = 3 biological replicates.

---

**Supplementary table S1:** Strains used in this study.

| Name | relevant genotype | Genotype | Source | Used in Figure(s) |
| --- | --- | --- | --- | --- |
| KWY165 | WT | *MATa, his3-11,15 ura3-1 leu2-3 trp1-1 ade2-1* | Joyner et al., 2016 | S2E, 4C, 4E, 4F, S4C |
| KWY8913 | GEMs | *MATa, his3-11,15 ura3-1 leu2-3 trp1-1 ade2-1 LEU2::pINO4-PfV-GS-Sapphire* | Kronig et al., 2026 | 1D, 1E, 1F, S1A, S1B, S1C, 3E, 3F, 3G, 3K, 3L, 3M, S3C, S3D, S3E, S3I, S3J, S3K |
| YGN2868 | TPKas | *MATa, his3-11,15 ura3-1 leu2-3 trp1-1 can1-100, GAL, psi+, ADE2, tpk1(M1646):NatNT2, tpk2(M1476):hphNT1, tpk3(M1656):His* | This Study | S2E |
| YGN2703 | TPKas | *MATalpha, leu2-3, ura3, trp1-1, his3-11,15, can1-100, GAL, psi+, ADE2 tpk1(M1646) tpk2(M1476) tpk3(M1656)* | This Study | 4C, 4E, 4F |
| YGN2782 | TPKas *msn2/4Δ* | *MATalpha, his3-11,15 ura3-1 leu2-3 trp1-1 can1-100, GAL, psi+, ADE2 tpk1(M1646) tpk2(M1476) tpk3(M1656) msn2::TRP1 msn4::KAN* | Kronig et al., 2026 (KWY11654) | 4C, 4F |
| YGN2781 | TPKas *glg1/2Δ glc3Δ* | *MATalpha, ade2-1, leu2-3, ura3, trp1-1, his3-11,15, can1-100, GAL, psi+, tpk1(M1646) tpk2(M1476) tpk3(M1656) TPKas, glg1::HygNT1 glg2::KanMX6 glc3::TRP* | This Study | 4C, 4F |
| KWY12340 | TPKas *gph1Δ* | *MATalpha, leu2-3, ura3, trp1-1, his3-11,15, can1-100, GAL, psi+, ADE2 tpk1(M1646) tpk2(M1476) tpk3(M1656) gph1Δ::URA3* | This Study | 4E |
| YGN3156 | TPKas *glc3Δ* | *MATa, leu2-3, ura3, trp1-1, his3-11,15, can1-100, GAL, psi+, ADE2 tpk1(M1646) tpk2(M1476) tpk3(M1656) TPKas, glc3::TRP* | This Study | 4F |
| YGN3391 | TPKas *glc3Δ tps1Δ* | *MATa, tpk1(M1646) tpk2(M1476) tpk3(M1656) TPKas, tps1::kanMX6 glc3::TRP* | This Study | 4F |
| YGN3685 | TPKas GEMs | *MATalpha, leu2-3, ura3, trp1-1, his3-11,15, can1-100, GAL, psi+, ADE2 tpk1(M1646) tpk2(M1476) tpk3(M1656) LEU2::pINO4-PfV-GS-Sapphire(pKW4635)* | This Study | 1D, 1E, 1F, S1A, S1B, S1C, 3E, 3F, 3G, 3K, 3L, 3M, S3C, S3D, S3E, S3I, S3J, S3K |
| KWY11785 | TPKas *msn2/4Δ* GEMs | *MATalpha, ade2-1, leu2-3, ura3, trp1-1, his3-11,15, can1-100, GAL, psi+, tpk1(M1646) tpk2(M1476) tpk3(M1656) TPKas, msn2::TRP1 msn4::KAN LEU2::pINO4-PfV-GS-Sapphire(pKW4635)* | This Study | 1D, 1E, 1F, S1A, S1B, S1C |
| KWY12078 | TPKas *glg1/2Δ glc3Δ* GEMs | *MATalpha, ade2-1, leu2-3, ura3, trp1-1, his3-11,15, can1-100, GAL, psi+, tpk1(M1646) tpk2(M1476) tpk3(M1656) glg1::HygNT1 glg2::KanMX6 glc3::TRP LEU2::pINO4-PfV-GS-Sapphire(pKW4635)* | This Study | 3E, 3F, 3G, S3C, S3D, S3E |
| YGN3715 | TPKas *tps1Δ* GEMs | *MATa, ade2-1, leu2-3, ura3, trp1-1, his3-11,15, can1-100, GAL, psi+, TPKas: tpk1(M1646):NatNT2, tpk2(M1476),* *tpk3(M1656):His, LEU2::pINO4-PfV-GS-Sapphire(pKW4635) (GEMs), tps1::Kan* | This Study | 3K, 3L, 3M, S3I, S3J, S3K |
| KWY9535 | *nth1Δ* | *MATa, his3-11,15 ura3-1 leu2-3 trp1-1 ade2-1 nth1Δ::HYG* | This Study | S4C |
| YGN2754 | *vph1-GFP* | *MATa, ade2-1, leu2-3, ura3, trp1-1, his3-11,15, can1-100, GAL, psi+, vph1-GFP:His3MX6* | This Study | 1A, 1B, 1C, 2D, 2E, S2A, S2B, S2D, 3A, 3H, S3A, S3F, S3H, 4A, 4B, S4A, S4B |
| YGN254 | *msn2/4Δ* | *MATa, ade2-1, leu2-3, ura3, trp1-1, his3-11,15, can1-100, GAL, psi+, msn2::TRP1, msn4::KAN* | This Study | S1D, S4A, S4B |
| YGN3107 | TPKas *vph1-GFP* | *MATa, ade2-1, leu2-3, ura3, trp1-1, his3-11,15, can1-100, GAL, psi+, TPKas: tpk1(M1646):NatNT2, tpk2(M1476):hphNT1, tpk3(M1656):His, vph1-GFP:TRP1* | This Study | 1A, 1B, 1C, 2A, 2B, 2C, 2D, 2E, S2A, S2D, 3A, 3B, 3C, 3D, 3H, 3I, 3J, S3H, 4A, 4B, 4D, 4G |
| YGN3153 | TPKas *glc3Δ vph1-GFP* | *MATalpha, leu2-3, ura3, trp1-1, his3-11,15, can1-100, GAL, psi+, ADE2 tpk1(M1646) tpk2(M1476) tpk3(M1656) TPKas, glc3::TRP, vph1-GFP:HIS3MX6* | This Study | 3A, 3B, 3C, 3D, 3H, 3I, 3J, S3H, 4A, 4B |
| YGN3189 | TPKas *msn2/4Δ vph1-GFP* | *MATa, ade2-1, leu2-3, ura3, trp1-1, his3-11,15, can1-100, GAL, psi+, tpk1(M1646) tpk2(M1476) tpk3(M1656) TPKas, msn2::TRP1, msn4::KAN, vph1-GFP:HIS3MX6* | This Study | 1A, 1B, 1C, 2E, S2A, 4A, 4B |
| YGN3261 | *gph1Δ* | *MATa, leu2-3,112 trp1-1 can1-100 ura3-1 ade2-1 his3-11,15 [phi+], gph1Δ::URA3* | This Study | S4A, S4B |
| YGN3404 | TPKas *gph1Δ* | *MATalpha, ade2-1, leu2-3, ura3, trp1-1, his3-11,15, can1-100, GAL, psi+, TPKas: tpk1(M1646):NatNT2, tpk2(M1476):hphNT1, tpk3(M1656):His, vph1-GFP:TRP1, gph1::URA 3* | This Study | 4A, 4B, 4D, 4G |
| YGN3435 | TPKas *nth1Δ* | *MATalpha, ade2-1, leu2-3, ura3, trp1-1, his3-11,15, can1-100, GAL, psi+, TPKas: tpk1(M1646):NatNT2, tpk2(M1476):hphNT1, tpk3(M1656):His, vph1-GFP:TRP1, nth1::kanMX6* | This Study | 4B |
| YGN3474 | *nth1Δ vph1-GFP* | *MATalpha, ade2-1, leu2-3,112, ura3, trp1-1, his3-11,15, can1-100, GAL, [psi+], nth1::TRP1, vph1-GFP:His3MX6* | This Study | S4B |
| YGN3529 | *glc3Δ vph1-GFP* | *MATa, ade2-1, leu2-3, ura3, trp1-1, his3-11,15, can1-100, GAL, psi+, vph1-GFP:His3MX6, glc3::TRP* | This Study | S3A, S3B, S3F, S3G, S4A, S4B |
| YGN3712 | TPKas *nth1Δ gph1Δ vph1-GFP* | *MATalpha, ade2-1, leu2-3, ura3, trp1-1, his3-11,15, can1-100, GAL, psi+, TPKas: tpk1(M1646):NatNT2, tpk2(M1476):hphNT1, tpk3(M1656):His, vph1-GFP:TRP1, nth1::kanMX6, gph1::URA* | This Study | 4B |
| YGN3393 | TPKas *tps1Δ glc3Δ vph1-GFP* | *MATalpha, leu2-3, ura3, trp1-1, his3-11,15, can1-100, GAL, psi+, ADE2, , tpk1(M1646) tpk2(M1476) tpk3(M1656) TPKas, glc3::TRP, tps1::kanMX6, Vph1-GFP:His3MX6* | This Study | 3H, 3I, 3J, S3H, 4B |
| YGN3399 | TPKas *tps1Δ vph1-GFP* | *MATa, leu2-3, ura3, trp1-1, his3-11,15, can1-100, GAL, psi+, ADE2, , tpk1(M1646) tpk2(M1476) tpk3(M1656) TPKas, tps1::kanMX6, Vph1-GFP:His3MX6* | This Study | 3H, 3I, 3J, S3H, 4B, 4F |
| YGN3531 | *tps1Δ vph1-GFP* | *MATalpha, ade2-1, leu2-3, ura3, trp1-1, his3-11,15, can1-100, GAL, psi+, vph1-GFP:His3MX6, tps1::kanMX6* | This Study | S3F, S3G, S4B |
| YGN3572 | *tps1Δ glc3Δ vph1-GFP* | *MATa, ade2-1, leu2-3, ura3, trp1-1, his3-11,15, can1-100, GAL, psi+, vph1-GFP:His3MX6, glc3::TRP, tps1::kanMX6* | This Study | S3F, S3G, S4B |
| YGN3164 | *URA3::CUP1-Msn2(S6A)-mKate2* | *MATa, leu2-3,112 trp1-1 can1-100 ura3-1 ade2-1 his3-11,15 [phi+], URA3::CUP1-Msn2(S6A)-mKate2* | This Study | S1E |
| YGN3522 | TPKas *vph1-GFP GSY2-mCherry* | *MATalpha, ade2-1, leu2-3, ura3, trp1-1, his3-11,15, can1-100, GAL, psi+, TPKas: tpk1(M1646):NatNT2, tpk2(M1476):hphNT1, tpk3(M1656):His, vph1-GFP:TRP1, GSY2-mCherry:Kann* | This Study | S2C |
| YGN3637 | TPKas *glc3Δ vph1-GFP GSY2-mCherry* | *MATalpha, leu2-3, ura3, trp1-1, his3-11,15, can1-100, GAL, psi+, ADE2 tpk1(M1646) tpk2(M1476) tpk3(M1656) TPKas, glc3::TRP, vph1-GFP:His3MX6, GSY2-mCherry:Kan* | This Study | S2C |
| YGN1103 | *Tom70-mCherry* | *MATa, ade2-1, ura3, trp1-1, his3-11,15, can1-100, GAL, psi+*<br>*Tom70-mCherry:KanMX*<br>*LEU2::pTPI1-neurosporaF0subunit9preseq-GFP-PEST-ADH1term::leu2* | This Study | S2E |
| YGN3033 | TPKas *Tom70-mCherry* | *MATa, leu2-3, ura3, trp1-1, his3-11,15, can1-100, GAL, psi+, ADE2, TPKas: tpk1(M1646):NatNT2, tpk2(M1476):hphNT1, tpk3(M1656):His, Tom70-mCherry:KanMX* | This Study | S2E |
| YGN2698 | *glg1/2Δ* | *Mata, leu2-3,112 trp1-1 can1-100 ura3-1 ade2-1 his3-11,15, glg1Δ::HygNT1 glg2Δ::KanMX6 [phi+]* | This Study | S3A |

**Supplementary Table S2:**
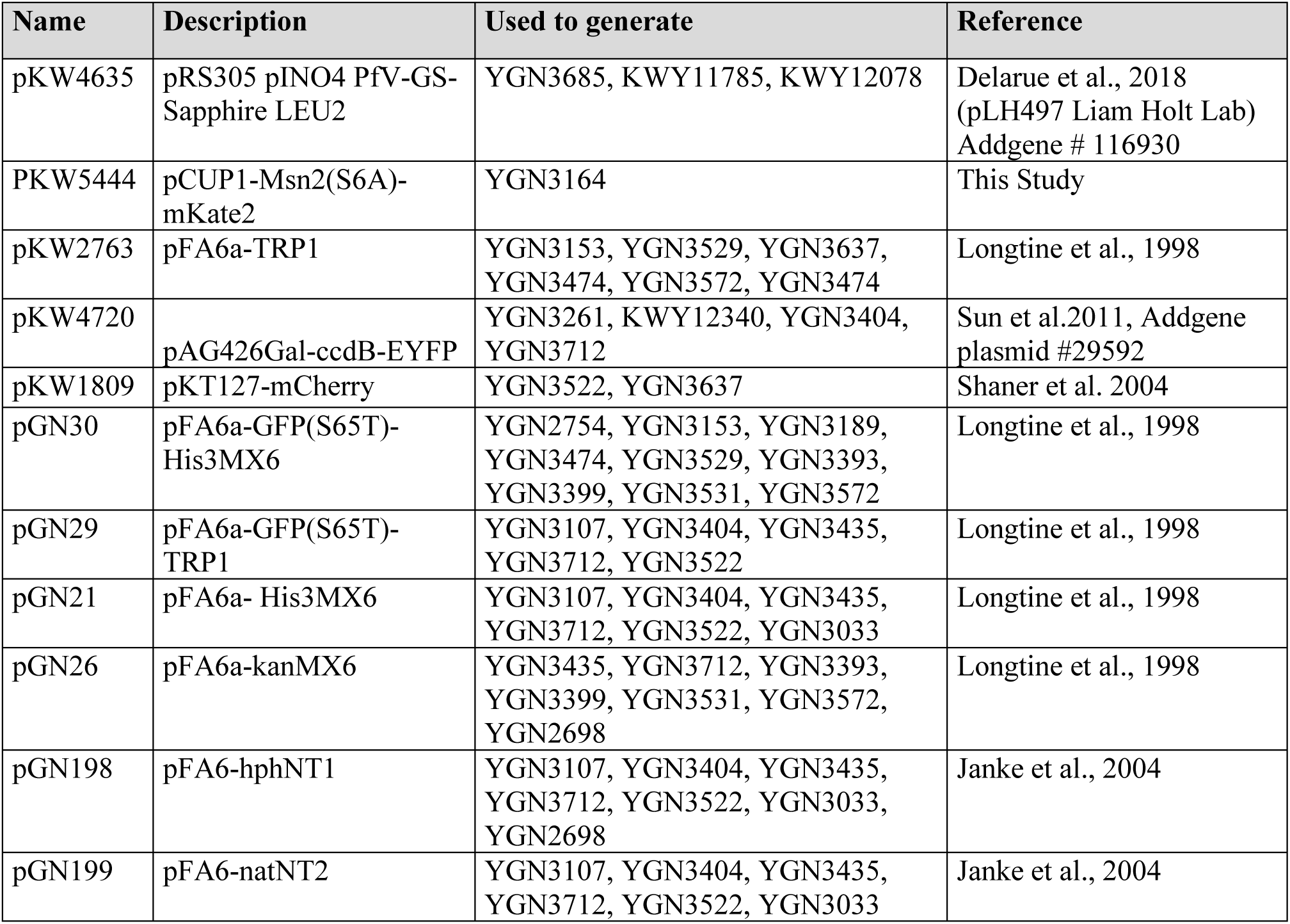
Plasmids.

**Supplementary table S3:**
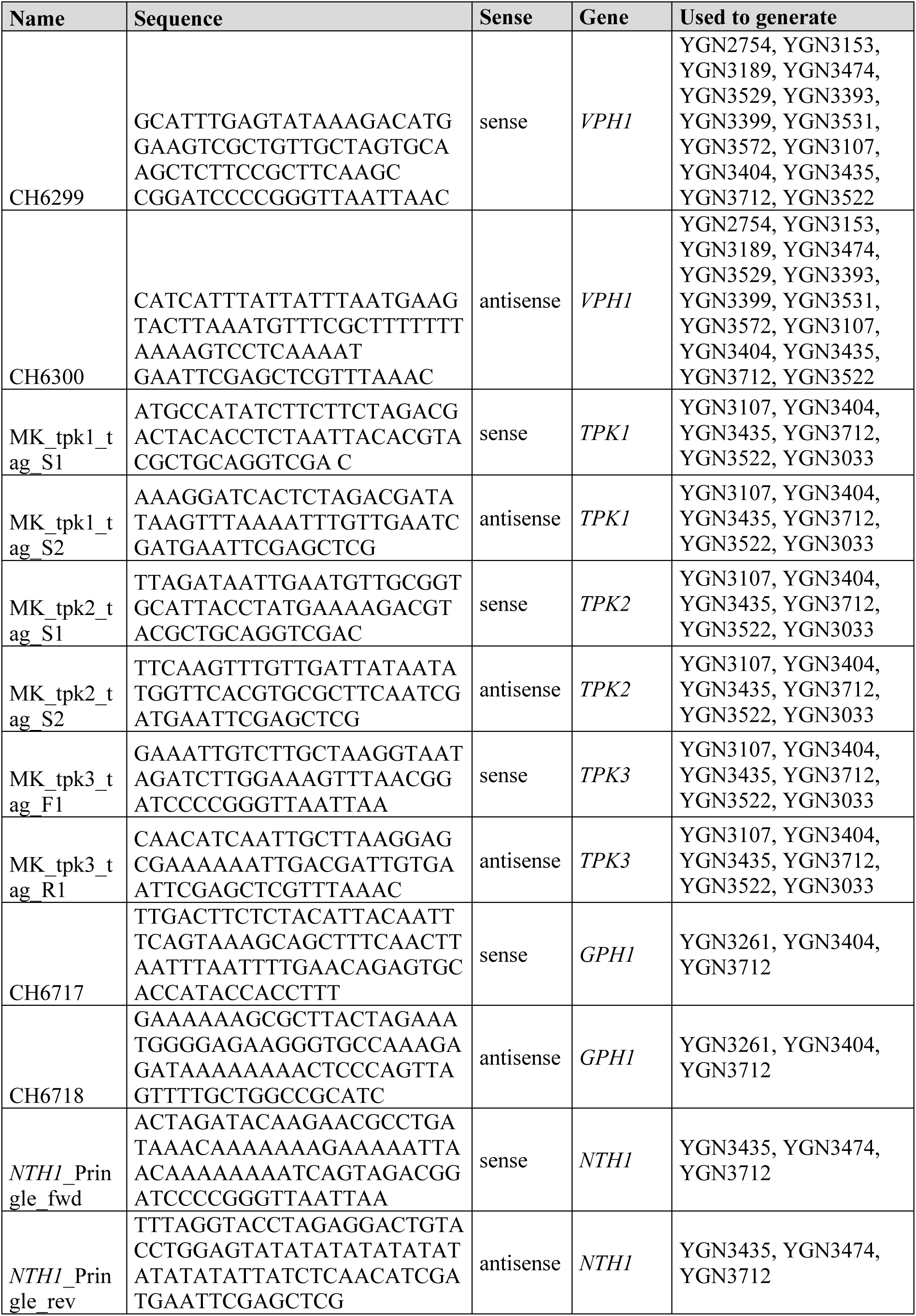
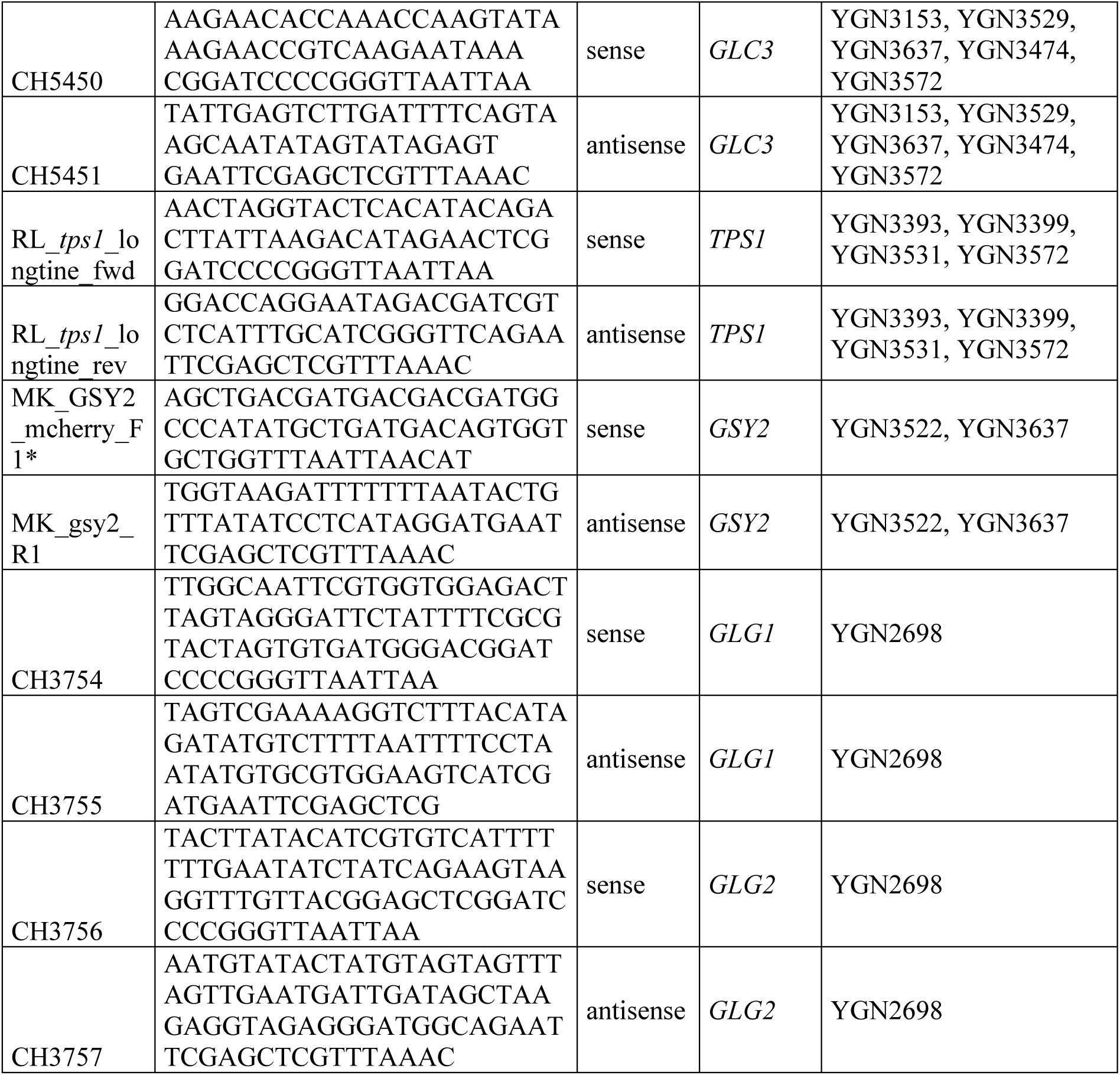
Primers.

## References

1. Pakos-Zebrucka, K., Koryga, I., Mnich, K., Ljujic, M., Samali, A., and Gorman, A.M. (2016). The integrated stress response. EMBO Rep. 17, 1374–1395. 10.15252/embr.201642195.

2. Berry, D.B., and Gasch, A.P. (2008). Stress-activated genomic expression changes serve a preparative role for impending stress in yeast. Mol. Biol. Cell 19, 4580–4587. 10.1091/mbc.e07-07-0680.

3. Dhar, R., Sägesser, R., Weikert, C., and Wagner, A. (2013). Yeast Adapts to a Changing Stressful Environment by Evolving Cross-Protection and Anticipatory Gene Regulation. Mol. Biol. Evol. 30, 573–588. 10.1093/molbev/mss253.

4. Gasch, A.P., Spellman, P.T., Kao, C.M., Carmel-Harel, O., Eisen, M.B., Storz, G., Botstein, D., and Brown, P.O. (2000). Genomic Expression Programs in the Response of Yeast Cells to Environmental Changes. Mol. Biol. Cell 11, 4241–4257. 10.1091/mbc.11.12.4241.

5. Causton, H.C., Ren, B., Koh, S.S., Harbison, C.T., Kanin, E., Jennings, E.G., Lee, T.I., True, H.L., Lander, E.S., and Young, R.A. (2001). Remodeling of yeast genome expression in response to environmental changes. Mol. Biol. Cell 12, 323–337. 10.1091/mbc.12.2.323.

6. Joyner, R.P., Tang, J.H., Helenius, J., Dultz, E., Brune, C., Holt, L.J., Huet, S., Müller, D.J., and Weis, K. (2016). A glucose-starvation response regulates the diffusion of macromolecules. eLife 5, e09376. 10.7554/eLife.09376.

7. Munder, M.C., Midtvedt, D., Franzmann, T., Nüske, E., Otto, O., Herbig, M., Ulbricht, E., Müller, P., Taubenberger, A., Maharana, S., et al. (2016). A pH-driven transition of the cytoplasm from a fluid- to a solid-like state promotes entry into dormancy. eLife 5:e09347. 10.7554/eLife.09347.

8. Delarue, M., Brittingham, G.P., Pfeffer, S., Surovtsev, I.V., Pinglay, S., Kennedy, K.J., Schaffer, M., Gutierrez, J.I., Sang, D., Poterewicz, G., et al. (2018). mTORC1 Controls Phase Separation and the Biophysical Properties of the Cytoplasm by Tuning Crowding. Cell 174, 338–349.e20. 10.1016/j.cell.2018.05.042.

9. Heimlicher, M.B., Bächler, M., Liu, M., Ibeneche-Nnewihe, C., Florin, E.-L., Hoenger, A., and Brunner, D. (2019). Reversible solidification of fission yeast cytoplasm after prolonged nutrient starvation. J. Cell Sci. 132, jcs231688. 10.1242/jcs.231688.

10. Neurohr, G.E., Terry, R.L., Lengefeld, J., Bonney, M., Brittingham, G.P., Moretto, F., Miettinen, T.P., Vaites, L.P., Soares, L.M., Paulo, J.A., et al. (2019). Excessive cell growth causes cytoplasm dilution and contributes to senescence. Cell 176, 1083–1097.e18. 10.1016/j.cell.2019.01.018.

11. Persson, L.B., Ambati, V.S., and Brandman, O. (2020). Cellular Control of Viscosity Counters Changes in Temperature and Energy Availability. Cell 183, 1572–1585.e16. 10.1016/j.cell.2020.10.017.

12. Klumpp, S., Scott, M., Pedersen, S., and Hwa, T. (2013). Molecular crowding limits translation and cell growth. Proc. Natl. Acad. Sci. U. S. A. 110, 16754–16759. 10.1073/pnas.1310377110.

13. Knapp, B.D., Odermatt, P., Rojas, E.R., Cheng, W., He, X., Huang, K.C., and Chang, F. (2019). Decoupling of Rates of Protein Synthesis from Cell Expansion Leads to Supergrowth. Cell Syst. 9, 434–445.e6. 10.1016/j.cels.2019.10.001.

14. Chen, Y., Huang, J.-H., Phong, C., and Ferrell, J.E. (2024). Viscosity-dependent control of protein synthesis and degradation. Nat. Commun. 15, 2149. 10.1038/s41467-024-46447-w.

15. Biswas, A., Muñoz, O., Kim, K., Hoege, C., Lorton, B.M., Nikolay, R., Kraushar, M.L., Shechter, D., Guck, J., Zaburdaev, V., et al. (2025). Conserved nucleocytoplasmic density homeostasis drives cellular organization across eukaryotes. Nat. Commun. 16, 7597. 10.1038/s41467-025-62605-0.

16. Xie, Y., Shu, T., Liu, T., Spindler, M.-C., Mahamid, J., Hocky, G.M., Gresham, D., and Holt, L.J. (2024). Polysome collapse and RNA condensation fluidize the cytoplasm. Mol. Cell 84, 2698–2716.e9. 10.1016/j.molcel.2024.06.024.

17. Gade, V.R., Heinrich, S., Paloni, M., Gómez-García, P.A., Dzanko, A., Oswald, A., Marchand, D., Khawaja, S., Barducci, A., and Weis, K. (2025). Polysomes and mRNA control the biophysical properties of the eukaryotic cytoplasm. Cell Rep. 44, 116204. 10.1016/j.celrep.2025.116204.

18. Kronig, L., Weber, C.A., Gómez-García, P.A., Fischer, J.S., Doerig, C., Michel, A., Ronchi, P., Khawaja, S., Picotti, P., Kornmann, B., et al. (2026). The environmental stress response regulates biophysics of the cytoplasm and survival in quiescence. J. Cell Biol. 225, e202601119. 10.1083/jcb.202601119.

19. Zaman, S., Lippman, S.I., Zhao, X., and Broach, J.R. (2008). How Saccharomyces Responds to Nutrients. Annu. Rev. Genet. 42, 27–81. 10.1146/annurev.genet.41.110306.130206.

20. Conrad, M., Schothorst, J., Kankipati, H.N., Van Zeebroeck, G., Rubio-Texeira, M., and Thevelein, J.M. (2014). Nutrient sensing and signaling in the yeast Saccharomyces cerevisiae. FEMS Microbiol. Rev. 38, 254–299. 10.1111/1574-6976.12065.

21. Creamer, D.R., Hubbard, S.J., Ashe, M.P., and Grant, C.M. (2022). Yeast Protein Kinase A Isoforms: A Means of Encoding Specificity in the Response to Diverse Stress Conditions? Biomolecules 12, 1–17. 10.3390/biom12070958.

22. Smith, A., Ward, M.P., and Garrett, S. (1998). Yeast PKA represses Msn2p/Msn4p-dependent gene expression to regulate growth, stress response and glycogen accumulation. EMBO J. 17, 3556–3564. 10.1093/emboj/17.13.3556.

23. Portela, P., Van Dijck, P., Thevelein, J.M., and Moreno, S. (2003). Activation state of protein kinase A as measured in permeabilised Saccharomyces cerevisiae cells correlates with PKA-controlled phenotypes in vivo. FEMS Yeast Res. 3, 119–126. 10.1016/S1567-1356(02)00158-7.

24. Wilson, W.A., Wang, Z., and Roach, P.J. (2005). Regulation of yeast glycogen phosphorylase by the cyclin-dependent protein kinase Pho85p. Biochem. Biophys. Res. Commun. 329, 161–167. 10.1016/j.bbrc.2005.01.106.

25. Schepers, W., Van Zeebroeck, G., Pinkse, M., Verhaert, P., and Thevelein, J.M. (2012). In Vivo Phosphorylation of Ser21 and Ser83 during Nutrient-induced Activation of the Yeast Protein Kinase A (PKA) Target Trehalase. J. Biol. Chem. 287, 44130–44142. 10.1074/jbc.M112.421503.

26. Shi, L., Sutter, B.M., Ye, X., and Tu, B.P. (2010). Trehalose is a key determinant of the quiescent metabolic state that fuels cell cycle progression upon return to growth. Mol. Biol. Cell 21, 1982– 1990. 10.1091/mbc.E10-01-0056.

27. Garay, E., Campos, S.E., González de la Cruz, J., Gaspar, A.P., Jinich, A., and DeLuna, A. (2014). High-Resolution Profiling of Stationary-Phase Survival Reveals Yeast Longevity Factors and Their Genetic Interactions. PLoS Genet. 10. 10.1371/journal.pgen.1004168.

28. De Virgilio, C., Bürckert, N., Bell, W., Jenö, P., Boller, T., and Wiemken, A. (1993). Disruption of TPS2, the gene encoding the 100-kDa subunit of the trehalose-6-phosphate synthase/phosphatase complex in Saccharomyces cerevisiae, causes accumulation of trehalose-6-phosphate and loss of trehalose-6-phosphate phosphatase activity. Eur. J. Biochem. 212, 315–323. 10.1111/j.1432-1033.1993.tb17664.x.

29. Hottiger, T., De Virgilio, C., Hall, M.N., Boller, T., and Wiemken, A. (1994). The role of trehalose synthesis for the acquisition of thermotolerance in yeast. II. Physiological concentrations of trehalose increase the thermal stability of proteins in vitro. Eur. J. Biochem. 219, 187–193. 10.1111/j.1432-1033.1994.tb19929.x.

30. Sola-Penna, M., and Meyer-Fernandes, J.R. (1998). Stabilization against Thermal Inactivation Promoted by Sugars on Enzyme Structure and Function: Why Is Trehalose More Effective Than Other Sugars? Arch. Biochem. Biophys. 360, 10–14. 10.1006/abbi.1998.0906.

31. Tapia, H., Young, L., Fox, D., Bertozzi, C.R., and Koshland, D. (2015). Increasing intracellular trehalose is sufficient to confer desiccation tolerance to Saccharomyces cerevisiae. Proc. Natl. Acad. Sci. U. S. A. 112, 6122–6127. 10.1073/pnas.1506415112.

32. Pepelnjak, M., Velten, B., Näpflin, N., Von Rosen, T., Palmiero, U.C., Ko, J.H., Maynard, H.D., Arosio, P., Weber-Ban, E., De Souza, N., et al. (2024). In situ analysis of osmolyte mechanisms of proteome thermal stabilization. Nat. Chem. Biol. 20, 1053–1065. 10.1038/s41589-024-01568-7.

33. Yorimitsu, T., Zaman, S., Broach, J.R., and Klionsky, D.J. (2007). Protein Kinase A and Sch9 Cooperatively Regulate Induction of Autophagy in Saccharomyces cerevisiae. Mol. Biol. Cell 18, 4180–4189. 10.1091/mbc.e07-05-0485.

34. Bishop, A.C., Ubersax, J.A., Petsch, D.T., Matheos, D.P., Gray, N.S., Blethrow, J., Shimizu, E., Tsien, J.Z., Schultz, P.G., Rose, M.D., et al. (2000). A chemical switch for inhibitor-sensitive alleles of any protein kinase. Nature 407, 395–401. 10.1038/35030148.

35. Lee, M., Kunzi, M., Neurohr, G., Lee, S.S., and Park, Y. (2023). Hybrid machine-learning framework for volumetric segmentation and quantification of vacuoles in individual yeast cells using holotomography. Biomed. Opt. Express 14, 4567–4578. 10.1364/BOE.498475.

36. Pfanzagl, V., Görner, W., Radolf, M., Parich, A., Schuhmacher, R., Strauss, J., Reiter, W., and Schüller, C. (2018). A constitutive active allele of the transcription factor Msn2 mimicking low PKA activity dictates metabolic remodeling in yeast. Mol. Biol. Cell 29, 2848–2862. 10.1091/mbc.E18-06-0389.

37. Parrou, J.L., Teste, M.-A., and François, J. (1997). Effects of various types of stress on the metabolism of reserve carbohydrates in Saccharomyces cerevisiae: genetic evidence for a stress-induced recycling of glycogen and trehalose. Microbiology 143, 1891–1900. 10.1099/00221287-143-6-1891.

38. Takeuchi, T., Iwamasa, T., and Miyayama, H. (1978). Ultrafine Structure of Glycogen Macromolecules in Mammalian Tissues. J. Electron Microsc. (Tokyo) 27, 31–38. 10.1093/oxfordjournals.jmicro.a050091.

39. Drochmans, P. (1962). Morphologie du glycogène: Etude au microscope électronique de colorations négatives du glycogène particulaire. J. Ultrastruct. Res. 6, 141–163. 10.1016/S0022-5320(62)90050-3.

40. Rowen, D.W., Meinke, M., and Laporte, D.C. (1992). GLC3 and GHAI of Saccharomyces cerevisiae Are Allelic and Encode the Glycogen Branching Enzyme. MOL CELL BIOL. 10.1128/mcb.12.1.22.

41. Thon, V.J., Vigneron-Lesens, C., Marianne-Pepin, T., Montreuil, J., Decq, A., Rachez, C., Ball, S.G., and Cannon, J.F. (1992). Coordinate regulation of glycogen metabolism in the yeast Saccharomyces cerevisiae. Induction of glycogen branching enzyme. J. Biol. Chem. 267, 15224– 15228. 10.1016/S0021-9258(18)42169-2.

42. Wilson, W.A., Boyer, M.P., Davis, K.D., Burke, M., and Roach, P.J. (2010). The subcellular localization of yeast glycogen synthase is dependent upon glycogen content. Can. J. Microbiol. 56, 408–420. 10.1139/w10-027.

43. Bharat, T.A.M., Hoffmann, P.C., and Kukulski, W. (2018). Correlative Microscopy of Vitreous Sections Provides Insights into BAR-Domain Organization In Situ. Structure 26, 879–886.e3. 10.1016/j.str.2018.03.015.

44. François, J., and Parrou, J.L. (2001). Reserve carbohydrates metabolism in the yeast *Saccharomyces cerevisiae*. FEMS Microbiol. Rev. 25, 125–145. 10.1111/j.1574-6976.2001.tb00574.x.

45. Deroover, S., Ghillebert, R., Broeckx, T., Winderickx, J., and Rolland, F. (2016). Trehalose-6-phosphate synthesis controls yeast gluconeogenesis downstream and independent of SNF1. FEMS Yeast Res. 16. 10.1093/femsyr/fow036.

46. Babazadeh, R., Lahtvee, P.-J., Adiels, C.B., Goksör, M., Nielsen, J.B., and Hohmann, S. (2017). The yeast osmostress response is carbon source dependent. Sci. Rep. 7, 990. 10.1038/s41598-017-01141-4.

47. Johansson, L., Virkki, L., Anttila, H., Esselström, H., Tuomainen, P., and Sontag-Strohm, T. (2006). Hydrolysis of β-glucan. Food Chem. 97, 71–79. 10.1016/j.foodchem.2005.03.031.

48. Weber, C.A., Sekar, K., Tang, J.H., Warmer, P., Sauer, U., and Weis, K. (2020). β-Oxidation and autophagy are critical energy providers during acute glucose depletion in Saccharomyces cerevisiae. Proc. Natl. Acad. Sci. U. S. A. 117, 12239–12248. 10.1073/pnas.1913370117.

49. Neurohr, G.E., and Amon, A. (2020). Relevance and regulation of cell density. Trends Cell Biol. 30, 213–225. 10.1016/j.tcb.2019.12.006.

50. Terhorst, A., Sandikci, A., Whittaker, C.A., Szórádi, T., Holt, L.J., Neurohr, G.E., and Amon, A. (2023). The environmental stress response regulates ribosome content in cell cycle-arrested S. cerevisiae. Front. Cell Dev. Biol. 11, 1118766. 10.3389/fcell.2023.1118766.

51. Bianco, E., Bonassera, M., Uliana, F., Tilma, J., Winkler, M., Zencir, S., Gossert, A., Oborská-Oplová, M., Dechant, R., Hugener, J., et al. (2025). Stm1 regulates Ifh1 activity revealing crosstalk between ribosome biogenesis and ribosome dormancy. Mol. Cell 85, 1806–1823.e17. 10.1016/j.molcel.2025.04.008.

52. Prats, C., Graham, T.E., and Shearer, J. (2018). The dynamic life of the glycogen granule. J. Biol. Chem. 293, 7089–7098. 10.1074/jbc.R117.802843.

53. Briza, P., Breitenbach, M., Ellinger, A., and Segall, J. (1990). Isolation of two developmentally regulated genes involved in spore wall maturation in Saccharomyces cerevisiae. Genes Dev. 4, 1775–1789. 10.1101/gad.4.10.1775.

54. Jiang, H., Song, C., Chen, C.-C., Xu, R., Raines, K.S., Fahimian, B.P., Lu, C.-H., Lee, T.-K., Nakashima, A., Urano, J., et al. (2010). Quantitative 3D imaging of whole, unstained cells by using X-ray diffraction microscopy. Proc. Natl. Acad. Sci. U. S. A. 107, 11234–11239. 10.1073/pnas.1000156107.

55. Sakai, K., Kondo, Y., Goto, Y., and Aoki, K. (2024). Cytoplasmic fluidization contributes to breaking spore dormancy in fission yeast. Proc. Natl. Acad. Sci. 121, e2405553121. 10.1073/pnas.2405553121.

56. Sakai, K., Kunzi, M., Uliana, F., Willig, L., Lappalainen, R., Kamada, Y., Neurohr, G.E., and Ikui, A.E. (2026). Cell wall remodeling is required for crowding homeostasis during cell growth in S. cerevisiae. Preprint at bioRxiv, 10.64898/2026.06.12.731398.

57. Huang, J.-H., and Ferrell, J.E. (2026). How does cytoplasmic crowding affect reaction rates? Mol. Cell 86, 9–23. 10.1016/j.molcel.2025.12.007.

58. Alric, B., Formosa-Dague, C., Dague, E., Holt, L.J., and Delarue, M. (2022). Macromolecular crowding limits growth under pressure. Nat. Phys. 18, 411–416. 10.1038/s41567-022-01506-1.

59. Srivastava, N., Calabrese, L., Plancke, C.N., Rollin, R., Venkova, L., Havas, K., Lagomarsino, M.C., and Piel, M. (2025). A Dual Homeostatic Regulation of Dry Mass and Volume Defines a Target Density in Proliferating Mammalian Cells. Preprint at bioRxiv, 10.1101/2025.04.24.650395.

60. Warner, J.R. (1999). The economics of ribosome biosynthesis in yeast. Trends Biochem. Sci. 24, 437–440. 10.1016/S0968-0004(99)01460-7.

61. Marenduzzo, D., Finan, K., and Cook, P.R. (2006). The depletion attraction: an underappreciated force driving cellular organization. J. Cell Biol. 175, 681–686. 10.1083/jcb.200609066.

62. Banani, S.F., Lee, H.O., Hyman, A.A., and Rosen, M.K. (2017). Biomolecular condensates: organizers of cellular biochemistry. Nat. Rev. Mol. Cell Biol. 18, 285–298. 10.1038/nrm.2017.7.

63. Pelletier, J.F., Field, C.M., Coughlin, M., Ryazanova, L., Sonnett, M., Wühr, M., and Mitchison, T.J. (2026). Glycogen-dependent demixing of frog egg cytoplasm at increased crowding. Biophys. J. 10.1016/j.bpj.2026.07.014.

64. Hatters, D.M., Minton, A.P., and Howlett, G.J. (2002). Macromolecular Crowding Accelerates Amyloid Formation by Human Apolipoprotein C-II *. J. Biol. Chem. 277, 7824–7830. 10.1074/jbc.M110429200.

65. Munishkina, L.A., Ahmad, A., Fink, A.L., and Uversky, V.N. (2008). Guiding Protein Aggregation with Macromolecular Crowding. Biochemistry 47, 8993–9006. 10.1021/bi8008399.

66. Gimón, I.V., Sandefur, C., and Schnell, S. (2026). Macromolecular crowding and protein aggregation: Friend, foe or contextual force? Prog. Biophys. Mol. Biol. 199, 79–98. 10.1016/j.pbiomolbio.2025.12.001.

67. Ajito, S., Hirai, M., Iwase, H., Shimizu, N., Igarashi, N., and Ohta, N. (2018). Protective action of trehalose and glucose on protein hydration shell clarified by using X-ray and neutron scattering. Phys. B Condens. Matter 551, 249–255. 10.1016/j.physb.2018.03.040.

68. Romero-Pérez, P.S., Moran, H.M., Cordone, D.P., Horani, A., Truong, A., Manriquez-Sandoval, E., Ramirez, J.F., Martinez, A., Gollub, E., Hunter, K., et al. (2025). Protein surface chemistry encodes an adaptive tolerance to desiccation. Cell Syst. 16. 10.1016/j.cels.2025.101407.

69. Jain, N.K., and Roy, I. (2009). Effect of trehalose on protein structure. Protein Sci. 18, 24–36. 10.1002/pro.3.

70. Longtine, M.S., Mckenzie Iii, A., Demarini, D.J., Shah, N.G., Wach, A., Brachat, A., Philippsen, P., and Pringle, J.R. (1998). Additional modules for versatile and economical PCR-based gene deletion and modification in Saccharomyces cerevisiae. Yeast 14, 953–961. 10.1002/(SICI)1097-0061(199807)14:10%3C953::AID-YEA293%3E3.0.CO;2-U.

71. Dunham, M., Gartenberg, M., and Brown, G.W. (2015). Methods in Yeast Genetics and Genomics, 2015 Edition: A CSHL Course Manual.

72. Schindelin, J., Arganda-Carreras, I., Frise, E., Kaynig, V., Longair, M., Pietzsch, T., Preibisch, S., Rueden, C., Saalfeld, S., Schmid, B., et al. (2012). Fiji: an open-source platform for biological-image analysis. Nat. Methods 9, 676–682. 10.1038/nmeth.2019.

73. Stirling, D.R., Swain-Bowden, M.J., Lucas, A.M., Carpenter, A.E., Cimini, B.A., and Goodman, A. (2021). CellProfiler 4: improvements in speed, utility and usability. BMC Bioinformatics 22, 433. 10.1186/s12859-021-04344-9.

74. Zhao, H., Brown, P.H., and Schuck, P. (2011). On the distribution of protein refractive index increments. Biophys. J. 100, 2309–2317. 10.1016/j.bpj.2011.03.004.

75. Ershov, D., Phan, M.-S., Pylvänäinen, J.W., Rigaud, S.U., Le Blanc, L., Charles-Orszag, A., Conway, J.R.W., Laine, R.F., Roy, N.H., Bonazzi, D., et al. (2022). TrackMate 7: integrating state-of-the-art segmentation algorithms into tracking pipelines. Nat. Methods 19, 829–832. 10.1038/s41592-022-01507-1.

76. Wieser, S., and Schütz, G.J. (2008). Tracking single molecules in the live cell plasma membrane—Do’s and Don’t’s. Methods 46, 131–140. 10.1016/j.ymeth.2008.06.010.

77. Resch, G.P., Brandstetter, M., Königsmaier, L., Urban, E., and Pickl-Herk, A.M. (2011). Immersion freezing of suspended particles and cells for cryo-electron microscopy. Cold Spring Harb. Protoc. 2011, 803–814. 10.1101/pdb.prot5642.

78. Haynes, R.M., Myers, J., López, C.S., Evans, J., Davulcu, O., and Yoshioka, C. (2025). A strategic approach for efficient cryo-EM grid optimization using design of experiments. J. Struct. Biol. 217, 108068. 10.1016/j.jsb.2024.108068.

79. Tivol, W.F., Briegel, A., and Jensen, G.J. (2008). An improved cryogen for plunge freezing. Microsc. Microanal. Off. J. Microsc. Soc. Am. Microbeam Anal. Soc. Microsc. Soc. Can. 14, 375–379. 10.1017/S1431927608080781.

80. Berger, C., Dumoux, M., Glen, T., Yee, N.B. -y, Mitchels, J.M., Patáková, Z., Darrow, M.C., Naismith, J.H., and Grange, M. (2023). Plasma FIB milling for the determination of structures in situ. Nat. Commun. 14, 629. 10.1038/s41467-023-36372-9.

81. Berger, C., Watson, H., Naismith, J.H., Dumoux, M., and Grange, M. (2025). Xenon plasma focused ion beam lamella fabrication on high-pressure frozen specimens for structural cell biology. Nat. Commun. 16, 2286. 10.1038/s41467-025-57493-3.

82. Mastronarde, D.N. (2005). Automated electron microscope tomography using robust prediction of specimen movements. J. Struct. Biol. 152, 36–51. 10.1016/j.jsb.2005.07.007.

83. Eisenstein, F., Fukuda, Y., and Danev, R. (2024). Smart parallel automated cryo-electron tomography. Nat. Methods 21, 1612–1615. 10.1038/s41592-024-02373-9.

84. Kremer, J.R., Mastronarde, D.N., and McIntosh, J.R. (1996). Computer visualization of three-dimensional image data using IMOD. J. Struct. Biol. 116, 71–76. 10.1006/jsbi.1996.0013.

85. Parrou, J.L., and François, J. (1997). A simplified procedure for a rapid and reliable assay of both glycogen and trehalose in whole yeast cells. Anal. Biochem. 248, 186–188. 10.1006/abio.1997.2138.

86. Quain, D.E. (1981). The determination of glycogen in yeasts. J. Inst. Brew. 87, 289–291. 10.1002/j.2050-0416.1981.tb04038.x.

